# Genome editing of papaya using both Cas9 and Cas12a

**DOI:** 10.1101/2025.05.22.655363

**Authors:** Jeremieh Hasley, Rey-Joseph Dinulong, Achyut Adhikari, David Christopher, Miaoying Tian

**Affiliations:** Department of Plant and Environmental Protection Sciences, University of Hawaii at Manoa, Honolulu, HI 96822; Department of Molecular Biosciences and Bioengineering, University of Hawaii at Manoa, Honolulu, HI 96822; Department of Plant Pathology, University of Georgia, Tifton, GA 31794

**Keywords:** CRISPR, Cas9, Cas12a, gene editing, papaya, *Carica papaya* L

## Abstract

Papaya (*Carica papaya* L.) is an economically important tropical crop that produces papain and highly nutritious fruit, which are used in the grocery, cosmetic, pharmaceutical, and food processing industries. However, various destructive pathogens severely threaten its production. Furthermore, limited natural genetic variation restricts breeding efforts for crop improvement. Therefore, we turned to gene editing as a tool to address these problems. We utilized two CRISPR systems (Cas9 and Cas12a) and two papaya genes, *CpPDS* (phytoene desaturase) and *CpMLO6* (Mildew Locus O 6), to establish efficient genome editing systems of papaya. The systems were delivered by an optimized protocol of *Agrobacterium*-mediated transformation (AMT) of embryogenic callus suspension cultures derived from hypocotyls. Accordingly, we transformed papaya with five plasmid constructs, each of which expressed one or two guide RNAs (gRNAs) for gene editing using either Cas9 or Cas12a. All except two T0 transgenic plants tested produced mutations with the majority containing indels of over 90%. Furthermore, successful mutation of the *CpPDS* gene using both Cas9 and Cas12a produced albino phenotypes as expected for disrupting a gene for carotenoid biosynthesis. Successful mutagenesis was achieved with seven out of eight gRNAs. Homozygous and/or biallelic mutants were generated from transformation using all five constructs, suggesting the feasibility of obtaining transgene-free homozygous segregating mutants by selfing in the second generation. Taken together, a robust and reliable papaya genome editing system was established, which enables genetic modification in various genomic environments to meet the diverse needs of basic scientific research and tropical crop improvement.

## Introduction

Adapted from bacterial immune systems, the clustered regularly interspaced short palindromic repeat (CRISPR)/CRISPR-associated protein (Cas) has revolutionized the way scientists perform functional genomics studies and crop breeding. The CRISPR technology has enabled and accelerated the functional genomics studies of numerous organisms, including many plant species (Ansori et al., 2023). It is a powerful tool for crop improvement as it can shorten breeding process by generating homozygous or complete knockout mutations in the first generation of transgenic plants and produce transgene-free plants either by segregating the transgene out in the following generation or through DNA-free gene editing (Navet and Tian, 2020b; Hasley et al., 2021; Bhattacharjee et al., 2023).

CRISPR/Cas9 system from *Streptococcus pyogenes* has been successfully used in genome editing of many plant species (Haque et al., 2018; Jaganathan et al., 2018). This gene editing system requires Cas9 and a single guide RNA (sgRNA), which is a fusion of CRISPR RNA (crRNA) containing a 20-nt DNA target sequence upstream of a Cas9 protospacer adjacent motif (PAM, 5’-NGG-3’) and trans-activating CRISPR RNA (tracrRNA). Guided by a sgRNA, the Cas9 nuclease cleaves the DNA target usually at 3-4 nt upstream of the PAM motif, generating a double-stranded break (DSB). In cells, the DSBs are repaired by non-homologous end joining (NHEJ), or homology-directed repair (HDR) when a homologous DNA template is provided, leading to random insertions and deletions (indels), or precise modifications guided by the DNA template, respectively. In addition to CRISPR/Cas9 system, the CRISPR from *Prevotella* and *Francisella* 1 (Cpf1, also known as Cas12a), is another tool widely used for plant gene editing (Endo et al., 2016; Wang et al., 2017; Xu et al., 2017; Jia et al., 2019; Li et al., 2019; Malzahn et al., 2019). Cas12a recognizes a T-rich PAM (TTTN or TTN) and has dual nuclease activity that cleaves the target DNA and processes its own crRNA (Wang et al., 2017; Bayat et al., 2018). It cleaves the target DNA strand 18-23 nt distal of the PAM, leaving staggering ends in contrast to blunt ends generated by Cas9 (Bandyopadhyay et al., 2020). These features allow editing of AT-rich regions of a genome that Cas9 is unable to access, convenient multiplex gene editing by expressing a single transcript containing multiple guide RNAs, and increased efficiency of precise editing via HDR. LbCpf1 (LbCas12a) from *Lachnospiraceae bacterium* ND2006 is broadly used for plant genome editing due to its highest activity in editing tests (Tang et al., 2017; Wang et al., 2017).

Papaya (*Carica papaya* L.) is an economically and scientifically important tropical fruit tree crop belonging to the family *Caricaceae*. Papaya fruit is highly nutritious (Ming et al., 2008) and consumed as a vegetable or fruit depending on the level of ripeness. Besides its use as food, the fruit, its enzyme extracts, and tree bark latex are widely used in the cosmetics, pharmaceutical and food processing industries (Ming et al., 2008; Carlos-Hilario and Christopher, 2015). The papaya plant is a large herbaceous perennial that can grow up to 30 feet. This diploid dicot has a relatively small genome of about 350 Mb (Ming et al., 2008; Yue et al., 2022), and a short life cycle, taking 4 months to flower and 9 months to produce ripe fruit after sowing (Aryal and Ming, 2014). It is genetically transformable with a reported regeneration time of 9-13 months (Fitch et al., 1993; Zhu et al., 2006). As such, papaya is considered an ideal model for the exploration of tropical-tree genomes and fruit-tree genomics (Ming et al., 2008). In addition, papaya is also a valuable and unique system for studying sex determination and differentiation in plants (Aryal and Ming, 2014; Liu et al., 2018). Papaya plants occur in one of the following three sex types: female, male and hermaphrodite. Sex determination in papaya has been attributed to sex chromosomes, but the sex determining gene(s) have yet to be defined (Aryal and Ming, 2014; VanBuren et al., 2015; Liu et al., 2018). One issue is that these genes reside in a non-recombining chromosomal region. Consequently, their identification, cloning and linkage to the sex phenotypes are impossible using traditional map-based cloning (Zhang et al., 2014). However, targeted mutagenesis through CRISPR genome editing holds the promise to genetically decipher the individual roles of these genes.

A diverse range of virus, fungal and oomycete pathogens hamper papaya production, including the papaya ringspot virus (PRSV), the oomycete, *Phytophthora palmivora* (Nelson, 2008), and the fungi, *Colletotrichum gloeosporioides* (Maeda and Nelson, 2014; Ong and Ali, 2015) and powdery mildews (Cunningham and Nelson, 2012; Vivas et al., 2017). PRSV is one of the most severe diseases of papaya, threatening papaya production worldwide (Gonsalves, 1998; Mishra et al., 2019). In the 1990s, it widely and rapidly spread through Hawaii’s major papaya growing areas and almost entirely wiped out the state’s papaya industry (Gonsalves, 1998). Fortunately, genetic engineering via coat-protein technology was successfully used to generate transgenic papaya with resistance against PRSV, which became the first commercialized transgenic fruit crop that saved Hawaii’s papaya industry (Gonsalves, 1998; Ferreira et al., 2002). However, creating resistance to the other diseases is not as straightforward because natural resistance and genetic variation against these pathogens are minimal, which makes disease resistance breeding through traditional crossing impossible and/or challenging. For example, intergeneric hybridization with papaya’s closest relative (*Vasconcellea goudotiana*) (Pereira et al., 2014) resulted in F1 sterility, and the few resulting progeny had low viability (Manshardt and Wenslaff, 1989; Hoover, 2016). Therefore, disease resistance breeding through advanced biotechnological approaches, including CRSPR, has the potential to safeguard papaya production.

It is apparent that a highly efficient CRISPR-mediated gene editing system for papaya is essential for functional genomics studies and breeding. Two previous studies reported the genome editing of papaya using CRISPR/Cas9. First, we recently used DNA-free genome editing via delivery of Cas9-sgRNA ribonucleoprotein complex to papaya protoplasts (Elias et al., 2023). Although we achieved efficient gene editing in protoplasts, regeneration of them to plants remains challenging. Another study (Brewer and Chambers, 2022) reported successful gene editing using the *Agrobacterium*-mediated transformation method described by Zhu et al. (2006). They targeted the papaya phytoene desaturase (*CpPDS*) gene; The mutations produced albino shoots (Brewer and Chambers, 2022). As is common, complete *PDS* knockouts cannot be fully regenerated through rooting (Brewer and Chambers, 2022). It is unclear how well this system would allow the regeneration of fully developed gene edited plants when targeting other genes for mutagenesis. The issue is the shoot regeneration media utilized in the transformation method, which was reported to cause rooting difficulty (Shimshock, 2016).

In the present study, we have streamlined the CRISPR-based knockout systems applied to papaya by simultaneously developing an optimized protocol for *Agrobacterium*-mediated transformation (AMT) and regeneration of whole plants. This was achieved by utilizing two CRISPR systems, Cas9 and Cas12a, to target two genes, *CpPDS* and *CpMLO6*. The latter encodes a homolog of Mildew Locus O (MLO), which is a potential susceptibility gene for creating powdery mildew resistance (Elias et al., 2023).

## Materials and Methods

### sgRNA selection and *in vitro* cleavage activity assay

Candidate guide RNA target sequences were identified using the EuPaGDT web server (http://grna.ctegd.uga.edu) with the mRNA sequences of *CpPDS* (NCBI Reference Sequence: XM_022033216.1) and *CpMLO6* (NCBI Reference Sequence: XM_022053717.1) as the queries on the SunUp Papaya genome assembly (Ming et al. 2008) (NCBI GenBank Assembly Accession: GCA_000150535.1) as a custom genome for on- and off-target analyses, as described (Elias et al., 2023). For Cas9 editing, 20-nt target sequences upstream of SpCas9 PAM motif (5’-NGG-3’) were identified. For Cas12a editing, 20-nt sequences downstream of LbCpf1 PAM motif (5’-TTTN-3’) were identified. Guide RNAs (gRNAs) with efficiency scores greater than 0.50 and no potential off-targets were subjected to secondary structure analyses using web tool, RNAstructure (http://rna.urmc.rochester.edu/RNAstructureWeb/Servers/Predict1/Predict1.html). Secondary structural analyses were conducted on Cas9 sgRNAs containing a 20-nt target sequence followed by a 76-nt scaffold sequence (Xing et al., 2014) and Cas12a crRNAs containing a 20-nt target sequence proceeded by a 21-nt direct repeat sequence (Wang et al., 2017). The gRNAs that maximally maintained the secondary structure of the Cas9 sgRNA scaffold sequence or the Cas12a direct repeat sequence, while simultaneously having minimal complexity in the target sequence, were selected for *in vitro* cleavage activity assays.

*In vitro* cleavage activities of sgRNAs for Cas9 and crRNAs for Cas12a were tested using the GenCrispr sgRNA Screening Kit (GenScript) and EnGen Lba Cas12a (New England Biolabs, NEB), respectively. The sgRNAs or crRNAs were synthesized by Integrated DNA Technologies (IDT). For each gRNA or crRNA, a PCR fragment of over 1 kb containing the target site was amplified using the corresponding primers (Table S1), Phusion high-fidelity DNA polymerase (NEB), and genomic DNA from cultivar Kapoho as a template. The cleavage assays of gRNAs for Cas9 were performed following the manufacturer’s instructions with or without 0.5 µl of 20 µM sgRNA in a 20-µl reaction. For *in vitro* cleavage activities of crRNAs for Cas12a, a similar protocol was used except that EnGen Lba Cas12a (NEB) and the corresponding reaction buffer were used in place of Cas9 and its buffer, and the temperature for incubating the crRNAs and Cas12a was 25 °C instead of 37 °C.

### Vector construction

For gene editing using Cas9, target sequences of two sgRNAs for each gene were cloned into pKSE401 (Xing et al., 2014; Navet and Tian, 2020b) to produce pKSE401::CpPDS and pKSE401::CpMLO6. For gene editing using Cas12a, we modified pKSE401 to pKSE401-Cas12a by replacing the sgRNA scaffold (sgRNA-Sc) and U6-26 terminator (U6-26t) with Poly T (TTTTTTTT), and zCas9 with LbCas12a. To achieve that, pKSE401(Xing et al., 2014) was digested using HindIII to remove (U6-26p)-(BsaI-SpR-BsaI)-(sgRNA-Sc)-(U6-26t), followed by the ligation of a HindIII-digested PCR fragment containing (U6-26p)-( BsaI-SpR-BsaI)-(PolyT) with HindIII restriction sites at both ends. The plasmid generated from the first step was named pKSE401-polyT. To replace zCas9 from pKSE401-polyT with Cas12a, XbaI and SacI restriction enzymes were used to remove zCas9. As the LbCas12a (Lbcpf1) sequence in pYPQ230 (Tang et al., 2017a) contains one XbaI and two SacI sites, an approach coupling overlapping PCR with mutagenesis was utilized to produce a LbCas12a fragment with synonymous mutations in the two SacI sites, and SpeI and SacI sites added at the two ends. This fragment was digested with SpeI and SacI, and then ligated into pKSE401-polyT with zCas9 removed with XbaI and SacI, to generate pKSE401-Cas12a. The sequence of pKSE401-Cas12a and a schematic map with key components are described in Supplemental File 1. For gene editing of *CpPDS* using Cas12a, the fragment containing full-length direct repeat (FLDR)-target 1-FLDR-target 2-FLDR (Xu et al. 2017; Wang et al. 2017), was cloned under the U6-26p promoter in pKSE401-Cas12a via two BsaI sites to construct pKSE401-Cas12a::CpPDS. For editing of *CpMLO6* using Cas12a, two plasmids with each expressing one crRNA were constructed by cloning FLDR-target-FLDR to pKSE401-Cas12a through two BsaI sites.

### Generation of liquid embryogenic cell cultures for papaya transformation

#### Seed germination

The *Carica papaya L.* cultivar, Kapoho, was used throughout this study. Seeds were soaked in a 1M KNO_3_ solution for 1 h with gentle shaking at 50 rpm at room temperature (RT). Soaked seeds were rinsed with deionized H_2_O (diH_2_O), and gently scrubbed in 1% liquinox using a short-bristled wire brush. After rinsing with diH_2_O, the seeds were surface sterilized in a solution containing 20% (v/v) Clorox Concentrated Germicidal Bleach (containing 8.25% sodium hypochlorite) and 40 µl/L of Tween-20 for 30 min with shaking at 100 rpm at RT. After rinsing 3-5 times with sterile MilliQ water, the seeds were sown in Magenta boxes containing seed germination media (1/2 MS [Murashige and Skoog Basal Medium], 0.5 g/L MES, pH 5.7 and 4.0 g/L Gelzan), and incubated at 29°C under a 12 h photoperiod with light intensity of 100 μmol m^−2^ s^−1^ until emergence of papaya seedlings.

#### Embryogenic callus induction

Hypocotyl tissues from the papaya seedlings, with fully expanded cotyledons and minimal growth of the first pair of true leaves, were cut into sections of 2 mm in length under aseptic conditions. The hypocotyl sections were placed onto callus induction media as described by (Carlos-Hilario and Christopher, 2015) with modifications, named mCIM10, which contained 1/2 MS, 30 g/L sucrose, 400 mg/L glutamine, 10 mg/L 2,4-D [2,4-Dichlorophenoxyacetic Acid], pH 5.7, and 3.2 g/L Gelzan, and incubated at 29°C in the dark for 5 to 8 weeks with bi-weekly subculturing until the production of friable embryogenic calli.

#### Establishment of papaya cell suspension culture

10-12 clumps of embryogenic calli at about 1.5 cm diameter, originating from different seedlings were collected and sliced into small pieces using a sterile sharp scalpel, and then placed into 50 ml of liquid callus induction media LCIM2 (1/2 MS, 60 g/L sucrose, 400 mg/L glutamine, 2 mg/L 2,4-D, pH 5.7) in a 250 ml Erlenmeyer flask. The culture was then incubated on a rotary shaker set to 120 rpm at 29°C in the dark for three weeks before it was ready for transformation. To maintain the cell suspension culture for later use, 2-3 ml of the established culture was transferred to 50 ml of fresh LCIM2 every 3 weeks.

### *Agrobacterium*-mediated transformation of papaya

*Agrobacterium tumefaciens* GV3101 containing the plasmids described above was used for transformation. Preparation of *Agrobacterium* was conducted as described previously (Navet and Tian, 2020a). An *Agrobacterium* suspension of OD_600_ = 0.4 in LCIM2 with 200 µM acetosyringone was prepared and incubated in the dark at room temperature for 1.5-2 h. Meanwhile, 50 ml of a 3-week-old papaya cell suspension culture grown in a flask was transferred to a 50 ml sterile tube and allowed to sit for 10 min, then 35 ml of liquid was removed. The remaining amount of the papaya cell suspension was divided to two portions, with each mixed with an equal volume of *Agrobacterium* expressing a gene editing construct, and incubated at RT in the dark with gentle sharking at 50 rpm for 30 min. The mixture was then centrifuged at 1000 g to pellet the cells. After removing most of the supernatant, the remaining liquid and cells (about 5 ml) were mixed and divided into five portions, with each transferred and evenly spread onto a piece of filter paper (Whatman 2) on top of CIM2 media (LCIM2, 3.2 g/L Gelzan) supplemented with 200 µM acetosyringone in a petri dish. After co-cultivation at 29°C in the dark for 2 days, the filter paper with *Agrobacterium* and papaya embryogenic calli was transferred to CIM2 media (LCIM2 with 3.2 g/L Gelzan) supplemented with 200 mg/L timentin, and incubated for 1 week at 29°C in the dark. The cells with the filter paper were then transferred and grown on CIM2 media supplemented with 150 mg/L timentin and 300 mg/L G418 (geneticin) for 6 weeks with subculturing every 3 weeks. Afterwards, the cells were scraped from the filter paper and placed in clusters onto embryo maturation media (CIM2 media without 2,4-D) supplemented with 100 mg/L timentin and 150 mg/L G418, and grown at 29°C in the dark for 3 weeks. For shoot regeneration, G418-resistant somatic embryos were grown on shoot regeneration media (1/2 MS, 20 g/L sucrose, 400 mg/L glutamine, 0.1 mg/L kinetin, 0.4 mg/L 6-BAP [6-Benzylaminopurine], pH 5.7, 3.2 g/L Gelzan) supplemented with 50 mg/L timentin and 150 mg/L G418, incubated at 29°C and a 12 h photoperiod with light intensity of 30-50 μmol m^−2^ s^−1^, and sub-cultured every three weeks for a course of 8 weeks to produce a sufficient number of elongated shoots. For root development, the elongated shoots were cultured in rooting medium (1/2 MS, 5 g/L sucrose, pH 5.7, 3.2 g/L Gelzan) with 50 mg/L timentin for 1 week, followed by 3 days of growing in rooting media with 0.5 mg/L IBA( Indole-3-butyric Acid), and then back on rooting medium supplemented with 50 mg/L timentin with subculturing every 3 weeks for about 3-8 weeks to allow a maximal number of shoots to develop roots. The same growth condition used for shoot regeneration was used for root development. Fully rooted plants were acclimatized and grown in soil as described (Navet and Tian, 2020a).

### Detection of mutations in transgenic plants

PCR fragments flanking the target sites were amplified using the primers listed in Table S1 and Phusion High-Fidelity DNA Polymerases (NEB), treated using ExoSAP-IT^TM^ (Thermo Fisher Scientific). Then they were sequenced by Sanger sequencing at Azenta Life Sciences. For detection of mutations, the sequencing chromatograms derived from transgenic plants and a wild-type (WT) plant were first analyzed using Sequencher (Gene Codes Corporation) to determine the quality of the sequencing and presence of the mutations. Sequencing chromatograms with mutations starting around the expected cleavage sites of Cas9 or Cas12a were decoded using Inference of CRISPR Editing (ICE) from Synthego, Inc. (https://ice.synthego.com/#/) to determine the type of insertions and deletions (indels).

## Results

### Identification of functionally active gRNAs

We identified two guide RNAs with *in vitro* cleavage activity for each target gene and Cas combination (Table 1). CpPDS-sg16 and CpPDS-sg379 were selected for Cas9-meditated gene editing of *CpPDS*. CpPDS-sg16 was expected to cleave a 1,256 bp amplicon generating two fragments of 968 bp and 288 bp, whereas CpPDS-sg379 was expected to cleave a 1,385 bp amplicon, generating fragments of 1,006 bp and 379 bp. Although the smaller bands of the cleaved products were faint on the gel due to the lower amount of DNA and the insensitivity of gel imager, the larger bands of the cleaved products were clearly seen (Fig. 1a). The intensity of non-cleaved bands was reduced in the reactions with both Cas9 and gRNAs compared with the controls in the absence of gRNAs, suggesting the functionality of sg16 and sg379 in the cleavage reaction (Fig. 1a). CpMLO6-sg254 and CpMLO6-sg439 were selected for gene editing of *CpMLO6* using Cas9. CpMLO6-sg254 cleaved a 1,250 bp amplicon producing fragments of 787 bp and 463 bp, while CpMLO6-sg439 cleaved a 1,328 bp amplicon into two fragments of 960 bp and 363 bp (Fig. 1b). CpPDS-cr253 and CpPDS-cr423rc were selected for Cas12-mediated gene editing of *CpPDS*, and were expected to cleave a 1,256 bp amplicon generating fragments at 663 bp and 593 bp for cr253, and to cleave a 1,385 bp amplicon generating fragments of 982 bp and 483 bp for cr423rc. For gene editing of *CpMLO6* using Cas12a, CpMLO6-cr292rc and CpMLO6-cr535 were identified and expected to cleave a 1,250 bp amplicon generating bands at 832 bp and 418 bp, and a 1,328 bp amplicon generating bands at 880 bp and 448 bp, respectively. For these four guide RNAs, the larger bands of the cleaved products were clearly visible on the agarose gel, suggesting that they are functional (Fig. 1c-d).

**Fig. 1.**
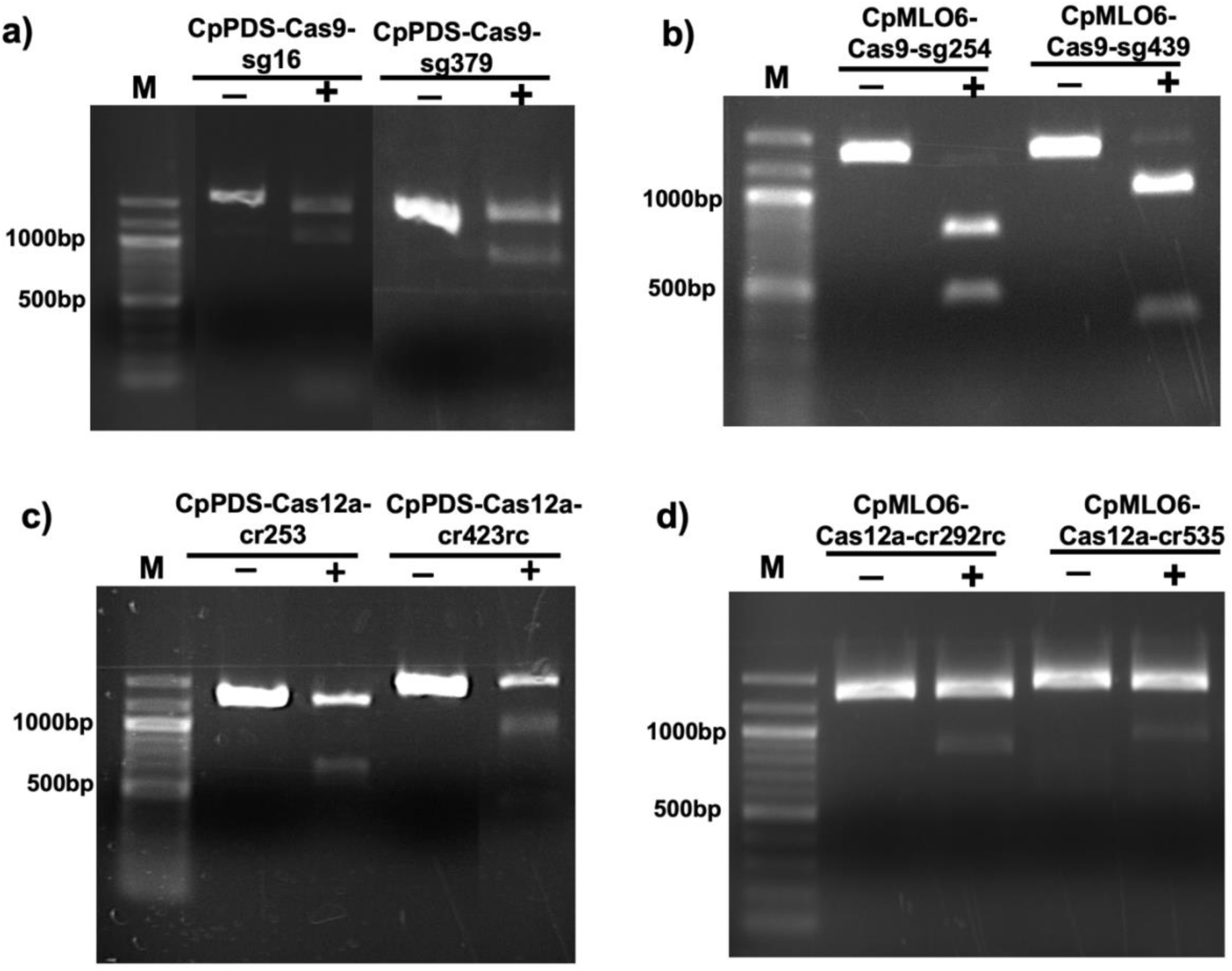
*In vitro* cleavage activities of gRNAs selected for gene editing of *CpPDS* and *CpMLO6* using Cas9 and Cas12a. For each gRNA indicated, the reaction contained the PCR fragment spanning its target sequence and Cas nuclease in the presence (+) or absence (-) of the gRNA. M, 100 bp DNA ladder (NEB).

**Table 1.**
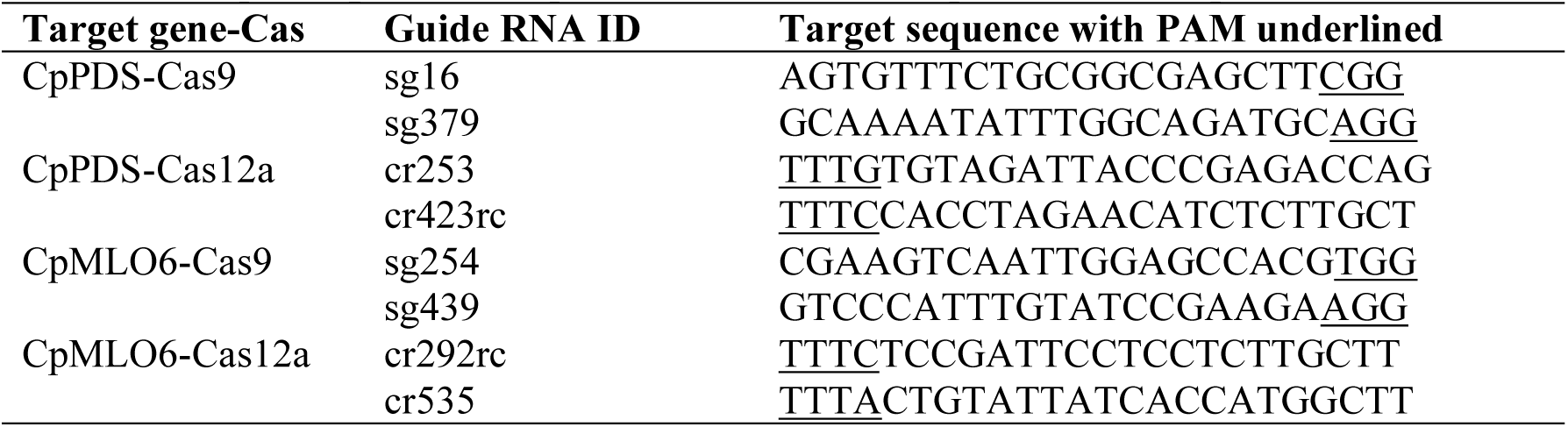
Target sequences of guide RNAs used for genome editing.

### Generation of constructs for papaya gene editing using Cas9 and Cas12a

For gene editing of *CpPDS* using Cas9, target sequences of sg379 and sg16 were cloned into pKSE401 to produce pKSE401::CpPDS with sg379 under the control of U6-26 promoter and sg16 under the control of U6-29 promoter, respectively (Fig. 2a). For gene editing of *CpMLO6* using Cas9, the target sequences of sg439 and sg254 were cloned into pKSE401 to produce pKSE401::CpMLO6 with the expression of sg439 driven by U6-26 promoter and the expression of sg254 driven by U6-29 promoter (Fig. 2b). pKSE401::CpPDS and pKSE401::CpMLO6 were constructed using the oligos in Table S2 following the methods as previously described (Xing et al., 2014; Navet and Tian, 2020b).

**Fig. 2.**
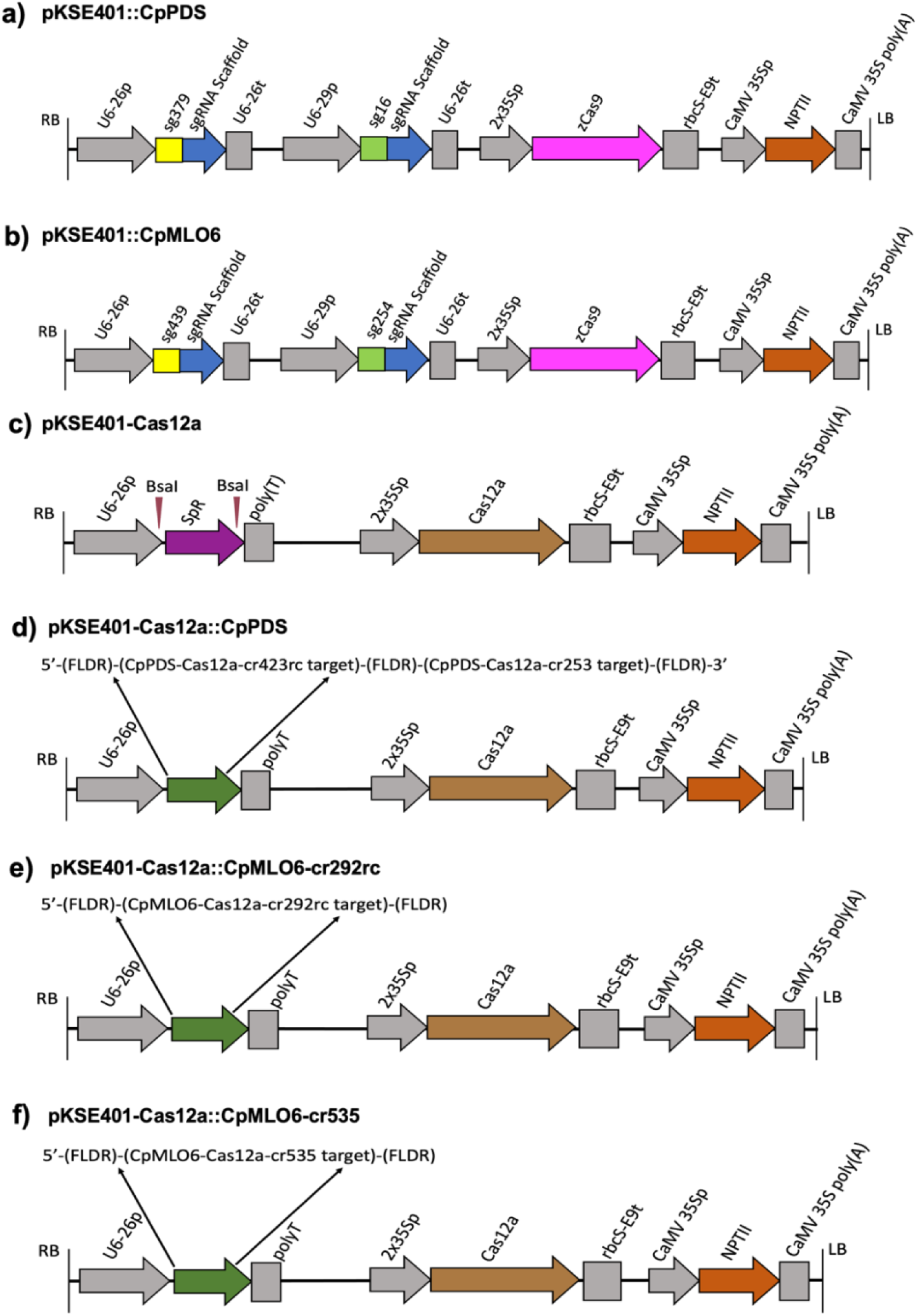
Schematic representations of expression cassettes within the T-DNA of the plasmids used for Cas9 editing of *CpPDS* (a) and *CpMLO6* (b), and Cas12a editing of *CpPDS* (d) and Cp*MLO6* (e-f) via *Agrobacterium*-mediated transformation. The vector modified from pKSE401 and used to clone crRNAs for Cas12 editing is shown in c). The labeling of the elements is as described by Xing et al. (2014).

For gene editing of *CpPDS* using Cas12a, the annealed oligos containing 35-nt full length direct repeat (FLDR), 20-nt cr423rc target sequence, FLDR, 20-nt cr253 target sequence, and then FLDR (Table S2), was cloned to pKSE401-Cas12a (Fig. 2c) to produce pKSE401-Cas12a::CpPDS, which expresses cr423rc and cr253 in an array under U6-26p promoter (Fig. 2d). For editing of *CpMLO6* using Cas12a, we constructed two plasmids pKSE401-Cas12a::CpMLO6-cr292rc (Fig. 2e) and pKSE401-Cas12a::CpMLO6-cr535 (Fig. 2f) with each expressing a single guide RNA cr292rc or cr535 to produce mutations in papaya through a single DNA break, a scenario in which the gene edited plants may be exempted from USDA regulation. These two plasmids were constructed by cloning the annealed oligos containing the 20-nt cr292rc or the cr535 target sequence, flanked by FLDR at each side (Table S2), into pKSE401-Cas12a.

### Optimization of *Agrobacterium*-mediated papaya transformation

We optimized a papaya transformation protocol based on a previously established *Agrobacterium*-mediated transformation method using embryogenic callus suspension cultures derived from hypocotyl tissues (Carlos-Hilario and Christopher, 2015) with significant modifications. Our detailed protocol utilized various medium components described previously for papaya and is described in the Materials and Methods. The workflows to produce embryogenic callus suspension cultures and to regenerate transgenic plants after infection of embryogenic calli with *Agrobacterium*, with representative images, are presented in Fig. 3 and Fig. 4, respectively. Here, we only highlight the significant modifications that were made.

**Fig. 3.**
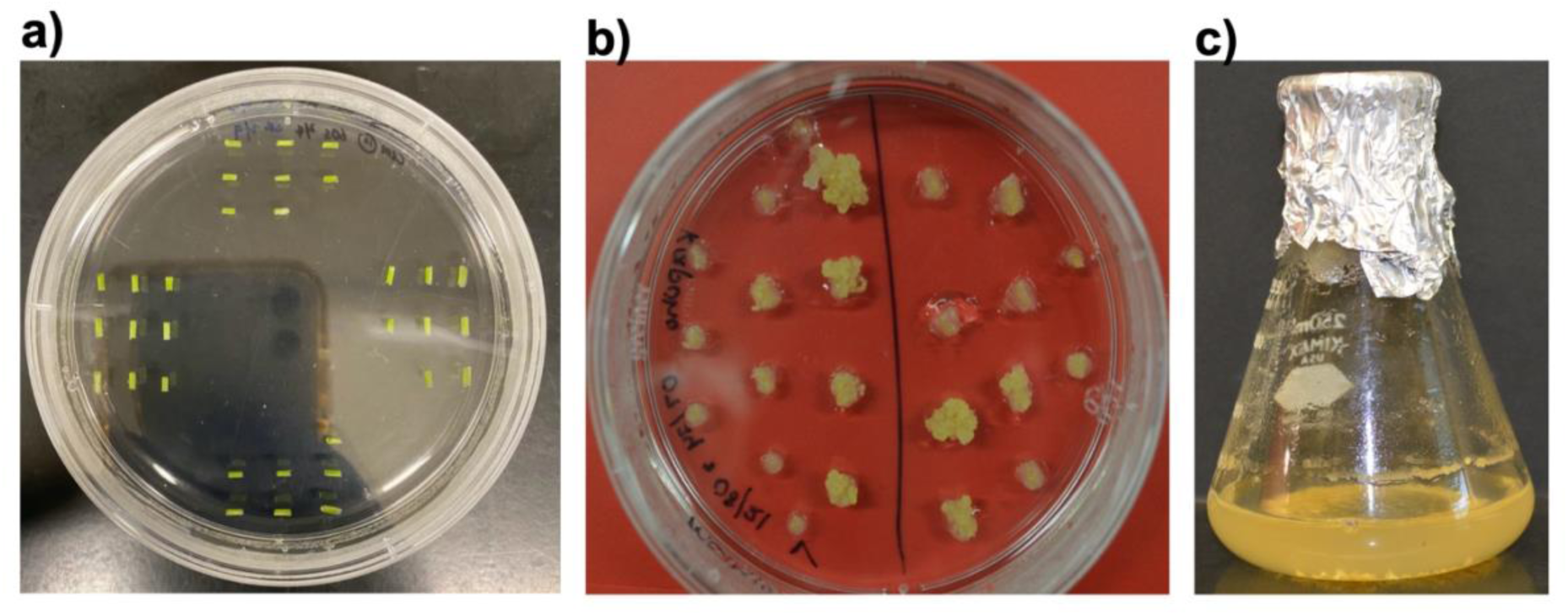
Establishment of hypocotyl-derived embryogenic calli cell suspension. a) 2 mm sections of papaya seedling hypocotyl tissue newly placed on mCIM10. b) Embryogenic calli proliferating on mCIM10 after 7 weeks. c) Embryogenic calli proliferating in LCIM2 after 3 weeks.

**Fig. 4.**
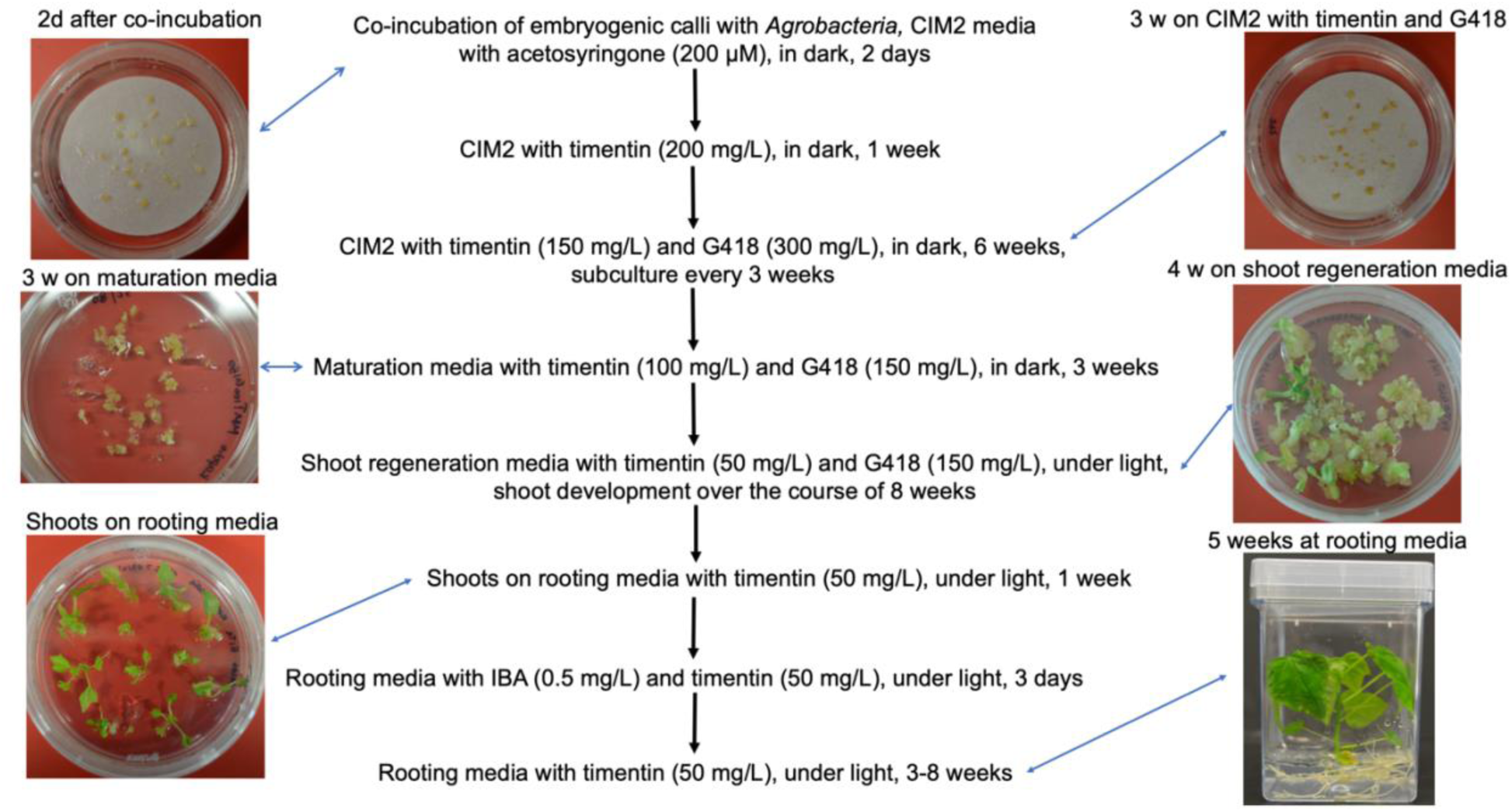
A workflow of selection and regeneration of papaya transgenic plants after *Agrobacterium*-mediated transformation of embryogenic calli. Representative photos at major stages of transformation using pKSE401::CpMLO6 are shown.

For induction of embryogenic calli, initially we used media that contained 70 g/L sucrose, as described (Zhu et al., 2006; Carlos-Hilario and Christopher, 2015), but only a low percentage of hypocotyl explants produced friable embryogenic calli after two months. After reducing the sucrose concentration to 30 g/L, a much higher percentage of explants produced embryogenic calli in a shorter time. We observed embryogenic calli at as early as 34 days after plating the hypocotyl tissues on mCIM10, and 118 out of 212 total explants produced embryogenic calli within 60 days.

For shoot regeneration from somatic embryos, instead of using MBN (MS medium supplemented with 2 mg/L each of 6-BAP and NAA [naphthalene acetic acid]) (Zhu et al., 2006; Carlos-Hilario and Christopher, 2015), we used the media containing 0.1 mg/L of kinetin and 0.4 mg/L of 6-BAP as described by (Ying et al., 1999) as shoots regenerated on MBN media were reported to be difficult to develop roots (Shimshock, 2016). For root regeneration, we utilized a rooting medium (1/2 MS, 5 g/L sucrose, pH 5.7, 3.2 g/L Gelzan) and protocol modified based on (Shimshock, 2016). As the high concentration of endogenous cytokinin affects root induction, we grew the shoots on the root medium without growth hormone for one week, and then on the medium with 0.5 mg/L of IBA for three days, followed by growing on the non-IBA rooting medium. Using this method, we were able to achieve about 50% of rooting efficiency when transforming pKSE401::CpMLO6 for editing *CpMLO6* using Cas9, which was higher than 26.3% achieved by (Shimshock, 2016). Similar rooting efficiency was achieved for shoots transformed with the constructs for editing *CpMLO6 via* Cas12a.

The constructs we used for this study express NPTII gene as a selectable marker (Fig. 2). Both kanamycin and G418 may be used for selection of transgenic somatic embryos and plants. It was reported that kanamycin significantly inhibited papaya somatic embryogenesis (Cai et al., 1999; Yu et al., 2003). Only after its removal, the kanamycin-resistant embryos were able to regenerate into shoots (Cai et al., 1999). To avoid that issue, we used G418 for selection, initially at 300 mg/L and reduced to 150 mg/L at the later steps until the regeneration of transgenic shoots (Fig. 4). For transformation with the five constructs, all regenerated shoots and/or plants, except two, produced mutations. We did not test whether these two lacking mutations were transformed. Nevertheless, this suggests that the selection scheme using G418 was successful in getting true transgenic plants.

To eliminate *Agrobacterium* after co-incubation with embryogenic calli, carbenicillin and/or cefotaxime were used previously with bacterial overgrowth issues (Fitch et al., 1993; Carlos-Hilario and Christopher, 2015). We used timentin initially at 200 mg/L and gradually reduced to 50 mg/L. No *Agrobacterium* overgrowth was observed. With this optimized protocol, we were able to obtain transgenic shoots for editing of *CpPDS* and plants for editing of *CpMLO6* using all five gene editing constructs. It took about 7 months to obtain fully regenerated gene edited transgenic plants from hypocotyl tissue explants, which is shorter than 9-13 months of regeneration time reported previously (Fitch et al., 1993; Zhu et al., 2006).

### Successful editing of the *CpPDS* gene using both Cas9 and Cas12a produced an albino shoot phenotype

Transformation of papaya using pKSE401::CpPDS (Fig. 2a) and pKSE401-Cas12a::CpPDS (Fig. 2d) regenerated albino shoots (Fig. 5), as predicted for mutagenizing a gene for the carotenoid pigment biosynthesis pathway. Carotenoids protect chlorophyll from photooxidative damage (Sun et al., 2018). As a result of the lack of chlorophyll, these shoots developed slowly and often formed cluster of leaves without a main stem. None of the albino shoots developed roots. We detected the mutations in seven albino plants transformed with pKSE401::CpPDS. All of them had an indel percentage greater than 97% at sg379 target site, suggesting a near-complete knockout of *CpPDS* in the T0 generation (Table 2). Two plants were homozygous with CpPDS-Cas9-1 having a 12nt deletion whereas CpPDS-Cas9-5 had a “T” insertion. The 12 nt deleted in CpPDS-Cas9-1 occurred at 3 nt upstream of the PAM motif (5’-AGG-3’) (Fig. 5a). All except the plant, CpPDS-Cas9-1, had short indels of 1nt insertions and 1-2 nt deletions. At the sg16 target site, good Sanger sequencing results were obtained from only two T0 plants (CpPDS-Cas9-6 and CpPDS-Cas9-7), both of which were chimeric and had an indel percentage of 98% containing a 1 nt insertion and a 4 nt deletion. These results suggest that high mutation efficiency was achieved by Cas9 and both gRNAs expressed under the U6-26 and U6-29 promoters, respectively.

**Fig. 5.**
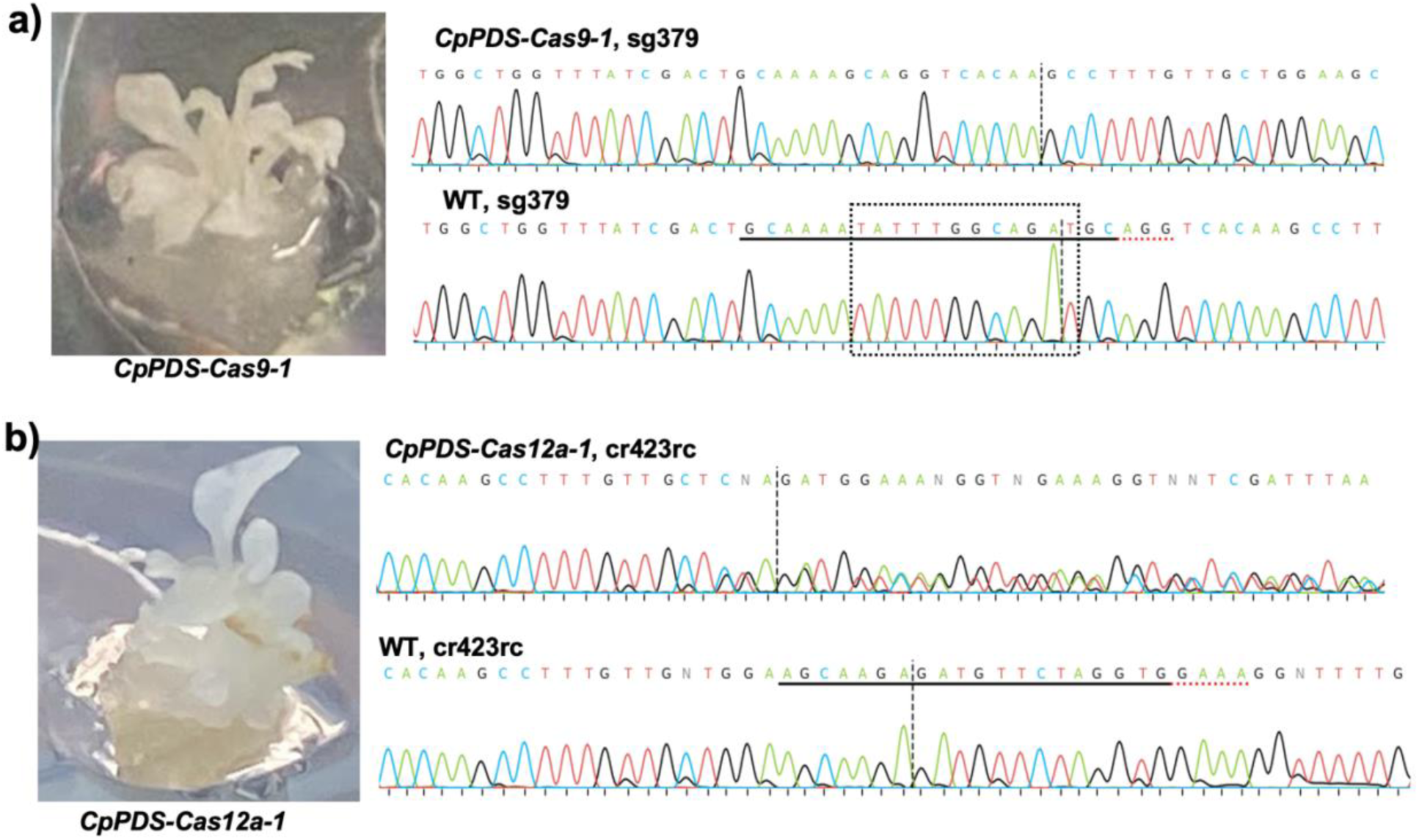
Gene editing of *CpPDS* produces albino shoots. Representative albino shoots generated by gene editing of *CpPDS* using Cas9 (a) or Cas12a (b), and their sequencing chromatograms at sg379 (a) or cr423rc (b) target site, respectively. Chromatograms of a wild-type plant (WT) at the corresponding target sites are shown. The target sequence of sg379 and the reverse complementary sequence of cr423rc target are underlined in the sequences of WT, with their PAM motifs underlined with red dotted lines. The 12 nt deleted in the mutant (CpPDS-cas9-1) are shown in a box with dotted lines.

**Table 2.**
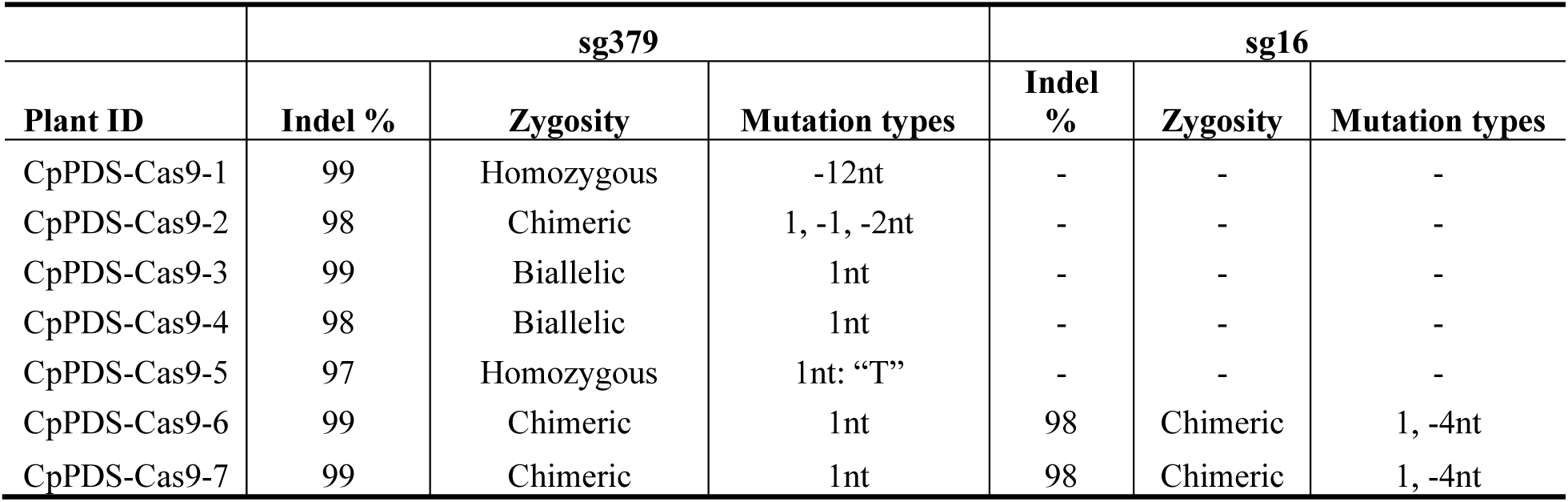
Mutations at sg379 and sg16 target sites in T0 pants transformed with pKSE401::CpPDS. nt, nucleotide. Under mutation types, a positive number indicates insertions, while a negative number indicates deletions. “-” indicates that the sequencing data were not available.

For Cas12-mediated editing of *CpPDS* by transforming with the plasmid pKSE401-Cas12a::CpPDS, all eight albino shoots had an indel percentage of over 90% at the cr423rc target site with two being biallelic and the remaining chimeric. A biallelic mutant plant with its sequencing chromatogram at cr423rc is shown in Fig. 5b. The mutation started at 23 nt distal to the PAM motif (5’-TTTC), with one allele having a 7nt deletion and another allele having a 16nt deletion based on the sequencing chromatogram decoding analysis using ICE (Synthego Co.). All mutations detected at the cr423rc target were deletions ranging from −4 nt to −16 nt. No mutations were detected at the cr253 target site in any of these plants.

### Editing of the *CpMLO6* locus using CRISPR/Cas9

We obtained over 40 transgenic plants regenerated from transformation using pKSE401::CpMLO6 (Fig. 2b). Mutations were tested in 26 of the transformants at both sg439 and sg254 target sites. Two T0 plants did not have mutations at either target site. The remaining 24 plants had mutations at both target sites (Table 3). At the sg439 target site, all 24 plants had mutation frequencies ranging from 93% to 100%. At the sg254 site, the mutation frequency ranged from 68% to 99%, with 20 plants having an indel percentage over 90%. At the sg439 site, the mutations were almost all short indels, including 1nt insertions and deletions of −1 nt to −6 nt. In addition to these short indels, the mutations at the sg254 sites included longer indels, such as 10-nt insertion and deletions of up to −35 nt. Homozygous, biallelic and chimeric mutations were detected at both target sites. At the sg439 site, 10 T0 plants contained homozygous mutations, including insertions of “A” or “T”, or deletion of −4 nt; eight contained biallelic mutations and six had chimeric mutations. At the sg254 target sites, the number of plants containing homozygous, biallelic and chimeric mutations was five, six, and thirteen, respectively. The homozygous mutations at the sg254 target site included a “C” insertion and deletions of −7 nt and −26 nt.

**Table 3.**
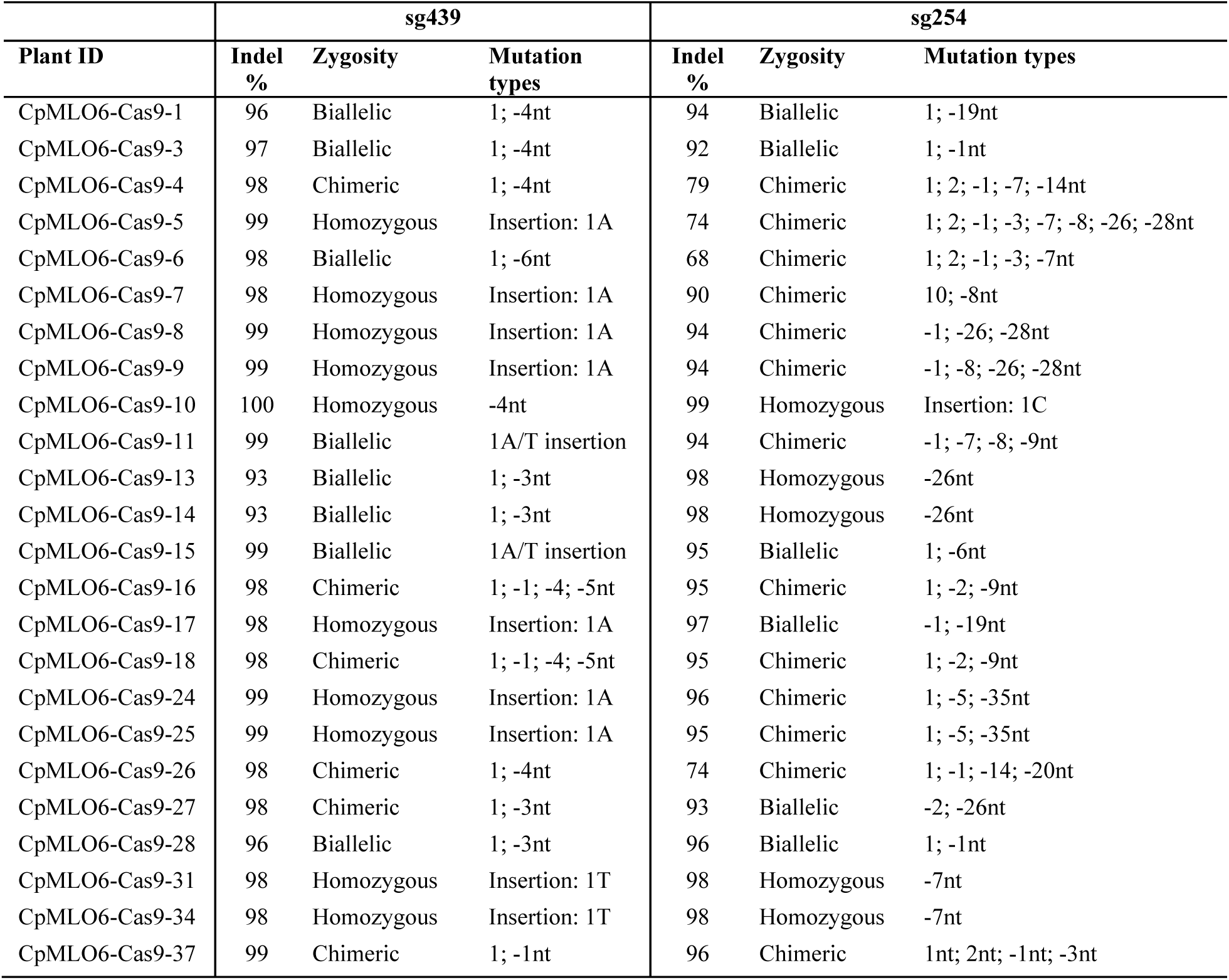
Mutations at sg439 and sg254 target sites in 24 T0 plants transformed with pKSE401::CpMLO6. nt, nucleotide. Under mutation types, a positive number indicates insertions, while a negative number indicates deletions.

### Editing of the *CpMLO6* locus using CRISPR/Cas12a

For editing of the *CpMLO6* locus using CRISPR/Cas12a, in order to generate mutations from a single DNA break, we made two constructs each expressing a single guide RNA, cr292rc or cr535 (Fig. 2e-f). We obtained 8 transgenic plants from transformation using pKSE401-Cas12a::CpMLO6-cr535 and 15 were regenerated from transformation using pKSE401-Cas12a::CpMLO6-cr292rc. For crRNA535, six out of eight plants contained mutations of over 89% frequency with one being homozygous for a 4 nt deletion, two biallelic and three chimeric; the remaining two were chimeric with an indel frequency of less than 45% (Fig. 6a). All mutations at cr535 were deletions ranging from −4 nt to −37 nt. The 4 nt deletion (5’-TGGC) in the homozygous plant cr535-CpMLO6-3 occurred at 15-18nt downstream of the PAM (5’-TTTA-3’) (Fig. 6c). For crRNA292rc, all transgenic plants were found to have biallelic mutations with over 90% indels frequency (Fig. 6b), with all mutations being deletions, ranging from −6 nt to −14 nt.

**Fig. 6.**
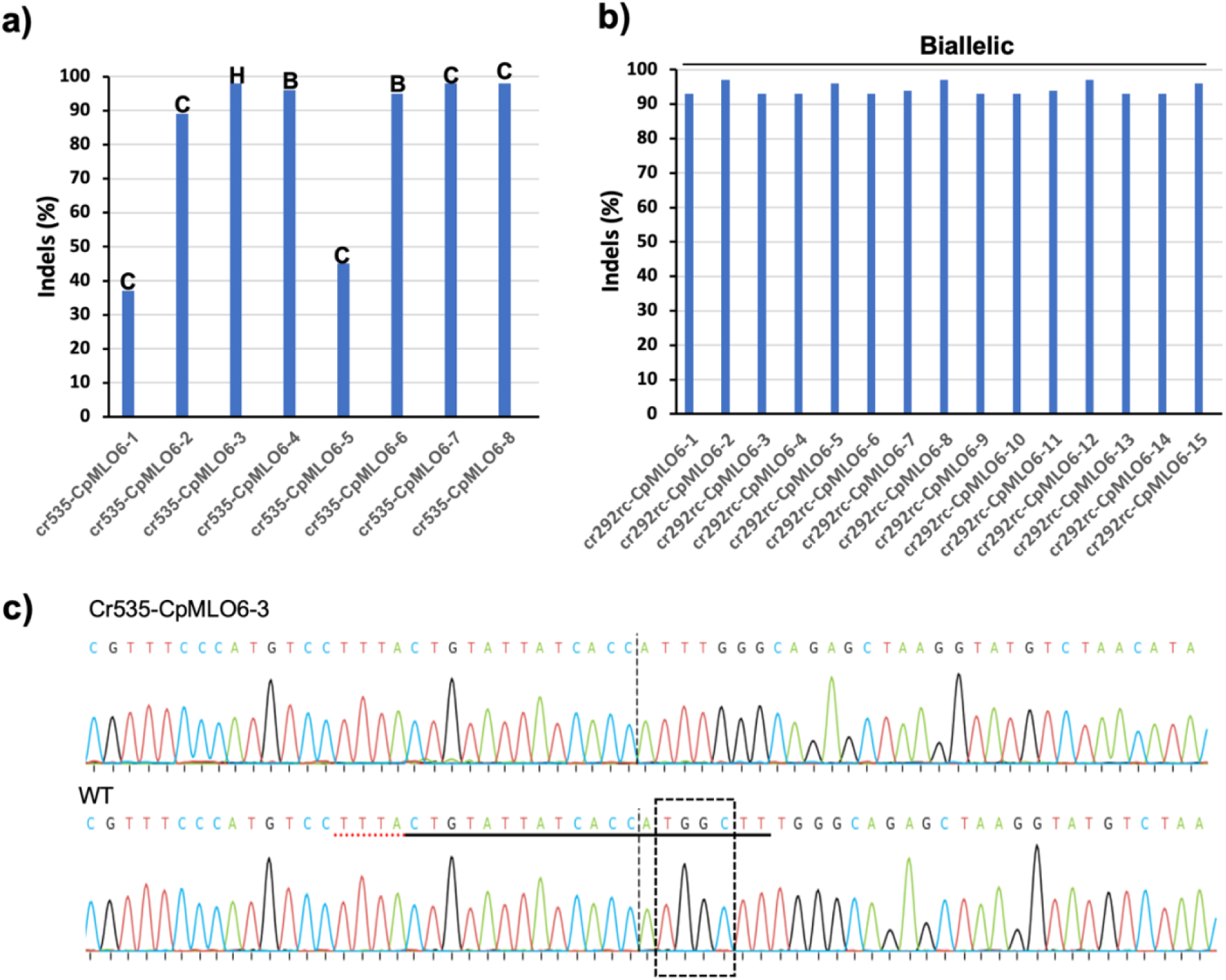
Editing of CpMLO6 using Cas12a. (a-b), Indel percentage of T0 plants expressing the crRNA535 (a) or cr292rc (b), with zygosity of the plants shown as H (homozygous), B (biallelic) and C (chimeric). c), Sequencing chromatograms of a wild-type plant (WT) and the homozygous mutant cr535-CpMLO6-3 at crRNA535 target site, with the target sequence underlined in solid line and PAM motif underlined with red dotted line, and the deleted 4 nucleotides shown in a box.

## Discussion

As papaya is a tropical fruit crop with agricultural, economical, and scientific significance, an efficient mutagenesis system is essential for its functional analysis, metabolic engineering and crop improvement. In this study, we established highly efficient papaya genome editing systems. This includes detailed protocols on: 1) the identification and *in vitro* cleavage activity assays of functional guide RNAs for both Cas9 and Cas12; 2) design and generation of an array of unique plasmid constructs for transformation; 3) establishment of papaya embryogenic callus cell suspension cultures from hypocotyls; 4) improved *Agrobacterium*-mediated transformation of these cultures, selection and regeneration of antibiotic-resistant transgenic plants; and 5) the detection of gene-specific mutations at the target sites. With these tools, we were able to successfully transform five constructs involving eight guide RNAs targeting eight sub-loci within two genes. This included two constructs for Cas9-mediated editing with each targeting two loci of either *CpPDS* or *CpMLO6*, one for Cas12a editing of *CpPDS* at two loci, and two for Cas12a editing of *CpMLO6* with each targeting a single locus. Except cr253, which was a guide RNA for genome editing of *CpPDS* using Cas12a, successful mutagenesis was achieved with all other seven guide RNAs, targeting seven genomic loci. All T0 transgenic plants tested for mutations, except two plants transformed with the plasmid pKSE401::CpMLO6), produced mutations with the majority of them containing indels of over 90%. Homozygous and/or biallelic mutants were generated from transformation of all constructs, suggesting the feasibility of producing transgene-free homozygous mutants in the next generation.

Cas9 was previously shown to be effective in papaya genome editing (Brewer and Chambers, 2022; Elias et al., 2023). In the present study, we expanded the gene targets for Cas9, and we demonstrated the effectiveness of Cas12a in genome editing of papaya, which extends the possible target loci to AT-rich regions that Cas9 is unable to access. The successful editing of the papaya genome using both Cas9 and Cas12a allows genetic modification at various genomic and diverse sequence environments to meet the needs of scientific research and breeding for desirable traits. For Cas9 editing, we utilized a binary vector, pKSE401, which is a vector previously created for gene editing in various dicots (Xing et al., 2014; Navet and Tian, 2020b; Hasley et al., 2021). As a similar vector for Cas12 editing was not readily available to use, we modified pKSE401 to pKSE401-Cas12a by replacing the CRISPR/Cas9 scaffold sequence and the U6-26 terminator (U6-26t) of pKSE401with Poly T, and zCas9 with LbCas12a. Using this newly modified vector, we successfully generated mutations in both *CpPDS1* and *CpMLO6*, suggesting the functionality of this vector. Like pKSE401, this vector is expected to serve as a useful workhorse for genome editing of papaya and other dicots.

We optimized the protocols for induction of embryogenic calli from papaya hypocotyls, *Agrobacterium*-mediated transformation of embryogenic callus cell suspensions, and the regeneration of transgenic plants. To achieve this, the media components from protocols in multiple previously published reports were modified and combined (details are presented in the Materials and Methods, and Results). The new protocols allowed us to generate embryogenic calli with a high frequency, improve rooting efficiency, overcome issues of *Agrobacterium* overgrowth and inhibition of somatic embryogenesis by kanamycin, to ultimately obtain transgenic plants in a time frame that is shorter than previously published methods. Using this system, we were able to successfully transform all five constructs tested, suggesting that this system is robust and reliable. Due to the handling of transformation with multiple constructs simultaneously, the amount of embryogenic calli we used for transformation with each construct was limited, and as such, the number of transgenic plants for some constructs was rather low. By increasing the amount of embryogenic calli used for infection by *Agrobacterium*, we expect to produce more transgenic plants with a higher percentage of homozygous mutants.

One of the advantages of the CRISPR/Cas12 is that a crRNA array containing multiple guide RNAs can be expressed under a single promoter, which is convenient for multiplex genome editing. For gene editing of *CpPDS* using Cas12a, a crRNA array containing a 35-nt full length direct repeat (FLDR), a 20-nt cr423rc target sequence, FLDR, a 20-nt cr253 target sequence, and then FLDR, was cloned under the U6-26 promoter, with the expectation of generating two functional guide RNAs that target two different loci. We observed indels of high percentage at the cr423rc target sites in all eight transgenic plants, but unexpectedly no mutations were detected at the cr253 target in any of these plants. It is unclear whether this is due to low editing efficacy of cr253, or an inability of processing the array to both mature cr423rc and cr253. Xu et al. (Xu et al., 2017) reported that expressing a pre-crRNA containing a 35-nt FLDR at either upstream or both ends of the target sequence resulted in increased mutation efficiency than expressing a mature crRNA with direct repeat (DR) followed by the target sequence. Our study clearly suggests that the use of the FLDR-target-FLDR is effective in generating mutations at a single target. Whether this strategy is effective for multiplex genome editing of papaya remains to be further investigated. Using the guide RNAs that have been shown to be effective in papaya editing, such as *CpMLO* cr292rc and cr535, or exchanging the order of cr253 and cr423rc in the crRNA array, is expected to provide the answer. Alternatively, we could test if a direct repeat (DR) instead of an FLDR functions in papaya multiplex gene editing as described in rice (Wang et al., 2017). Nevertheless, the option of multiplex gene editing in papaya is available by using Cas9. For transforming pKSE401::CpPDS and pKSE401::CpMLO6, mutations were detected at both sgRNA target sites of each gene. As the toolkit including pKSE401 as a component allows the cloning of two or more sgRNAs, it is entirely feasible to perform multiplex gene editing in papaya when necessary.

## Acknowledgments

We sincerely thank Dr. Qi-Jun Chen from China Agricultural University for providing the plasmid pKSE401 (Addgene Plasmid #62202), and Dr. Yiping Qi of University of Maryland for the plasmid pYPQ230 (Addgene plasmid #86210).

## Statements and Declarations

### Author Contributions

Conceptualization: MT, DC; Formal analysis and investigation: JH, RJD, MT, AA; Writing-original draft preparation: MT, JH; Writing – review & editing: MT, DC; Supervision: MT, AA; Funding acquisition: MT, DC. All authors read and approved the final manuscript.

## Supplementary information

**Supplemental File 1. Sequence and schematic map of pKSE401-Cas12a**

**Table S1. Target sequences of guide RNAs and primers used to amplify the fragments for *in vitro* cleavage activity assays and mutation detection**

**Table S2. Oligos used for constructing the plasmids for genome editing of *CpPDS* and *CpMLO6*.** The sequences and reverse complementary sequences of the targets of guide RNAs are underlined. The full-length direct repeats (FLDR) of CRISPR-Cas12a are shaded.

