## Supplemental File 1 for "Genome editing of papaya using both Cas9 and Cas12a"

**Supplemental File 1. Sequence and schematic map of pKSE401-Cas12a**

**I. pKSE401-Cas12a full sequence (15,693bp) with key elements related to the plasmid modification and guide RNA cloning highlighted.**

HindIII (bold, underlined), U6-26p (light blue), BsaI (upper case, bold and underlined), SpR (green), BsaI (upper case, bold and underlined), HindIII (bold, underlined), polyT (dark blue), 2x35S Promoter (Yellow), LbCpf1(LbCas12a)(Magenta) with two synonymous mutations from the sequence in pYPQ230 (Tang et al. 2017) shown (bold, italic and underlined).

taaacgctcttttctcttaggtttacccgccaatatatcctgtcaaacactgatagtttaaactgaaggcgggaaacgacaatctgatccaagctcaagctgctctagcattcgccattcaggctgcgcaactgttgggaagggcgatcggtgcgggcctcttcgctattacgccagctggcgaaagggggatgtgctgcaaggcgattaagttgggtaacgccagggttttcccagtcacgacgttgtaaaacgacggccagtgcc***aagctt***cgacttgccttccgcacaatacatcatttcttcttagctttttttcttcttcttcgttcatacagtttttttttgtttatcagcttacattttcttgaaccgtagctttcgttttcttctttttaactttccattcggagtttttgtatcttgtttcatagtttgtcccaggattagaatgattaggcatcgaaccttcaagaatttgattgaataaaacatcttcattcttaagatatgaagataatcttcaaaaggcccctgggaatctgaaagaagagaagcaggcccatttatatgggaaagaacaatagtatttcttatataggcccatttaagttgaaaacaatcttcaaaagtcccacatcgcttagataagaaaacgaagctgagtttatatacagctagagtcgaagtagtgattgg**GAGACC**aacccagtggacataagcctgttcggttcgtaagctgtaatgcaagtagcgtatgcgctcacgcaactggtccagaaccttgaccgaacgcagcggtggtaacggcgcagtggcggttttcatggcttgttatgactgtttttttggggtacagtctatgcctcgggcatccaagcagcaagcgcgttacgccgtgggtcgatgtttgatgttatggagcagcaacgatgttacgcagcagggcagtcgccctaaaacaaagttaaacatcatgggggaagcggtgatcgccgaagtatcgactcaactatcagaggtagttggcgtcatcgagcgccatctcgaaccgacgttgctggccgtacatttgtacggctccgcagtggatggcggcctgaagccacacagtgatattgatttgctggttacggtgaccgtaaggcttgatgaaacaacgcggcgagctttgatcaacgaccttttggaaacttcggcttcccctggagagagcgagattctccgcgctgtagaagtcaccattgttgtgcacgacgacatcattccgtggcgttatccagctaagcgcgaactgcaatttggagaatggcagcgcaatgacattcttgcaggtatcttcgagccagccacgatcgacattgatctggctatcttgctgacaaaagcaagagaacatagcgttgccttggtaggtccagcggcggaggaactctttgatccggttcctgaacaggatctatttgaggcgctaaatgaaaccttaacgctatggaactcgccgcccgactgggctggcgatgagcgaaatgtagtgcttacgttgtcccgcatttggtacagcgcagtaaccggcaaaatcgcgccgaaggatgtcgctgccgactgggcaatggagcgcctgccggcccagtatcagcccgtcatacttg^2^aagctagacaggcttatcttggacaagaagaagatcgcttggcctcgcgcgcagatcagttggaagaatttgtccactacgtgaaaggcgagatcaccaaggtagtcggcaaataatgtctagctagaaattcgttcaagccgacgccgcttcgcggcgcggcttaactcaagcgttagatgcactaagcacataattgctcacagccaaactatcaggtcaagtctgcttttattatttttaagcgtgcataataagcc**GGTCTC**gtttttttt***aagctt***gcatgcctgcaggtcaacatggtggagcacgacacacttgtctactccaaaaatatcaaagatacagtctcagaagaccaaagggcaattgagacttttcaacaaagggtaatatccggaaacctcctcggattccattgcccagctatctgtcactttattgtgaagatagtggaaaaggaaggtggctcctacaaatgccatcattgcgataaaggaaaggccatcgttgaagatgcctctgccgacagtggtcccaaagatggacccccacccacgaggagcatcgtggaaaaagaagacgttccaaccacgtcttcaaagcaagtggattgatgtgataacatggtggagcacgacacacttgtctactccaaaaatatcaaagatacagtctcagaagaccaaagggcaattgagacttttcaacaaagggtaatatccggaaacctcctcggattccattgcccagctatctgtcactttattgtgaagatagtggaaaaggaaggtggctcctacaaatgccatcattgcgataaaggaaaggccatcgttgaagatgcctctgccgacagtggtcccaaagatggacccccacccacgaggagcatcgtggaaaaagaagacgttccaaccacgtcttcaaagcaagtggattgatgtgatatctccactgacgtaagggatgacgcacaatcccactatccttcgcaagacccttcctctatataaggaagttcatttcatttggagaggacctcgacctcaacacaacatatacaaaacaaacgaatctcaagcaatcaagcattctacttctattgcagcaatttaaatcatttcttttaaagcaaaagcaattttctgaaaattttcaccatttacgaacgatactcgagtaaTCTAGtatggctcctaagaagaagcggaaggttggtattcacggggtgcctgcggcttcaaagctcgagaaattcaccaactgttattcgttgagcaaaacactgcggtttaaagcgattccagtcggcaagactcaagagaatatagacaataagcggctgttggtggaagatgaaaagcgcgcggaagactacaaaggggtgaagaagttgttggacagatactacctctcttttatcaatgatgtcttgcactcaatcaaattgaagaatctgaacaactacatctccctcttcagaaagaaaacaaggacagaaaaggagaataaggaacttgaaaatttggagatcaatctgaggaaagagatcgcgaaagcctttaaaggcaacgaaggatacaaaagtctgttcaagaaggatataattgagacaattttgccagagttcctcgatgacaaggacgagattgcgctggtcaattcgttcaacggattcacaacagcattcacaggcttctttgataatcgggaaaatatgttctctgaggaggcaaagtccacttctattgcgttcaggtgtatcaatgagaatctcactaggtacatttccaacatggatatctttgagaaggttgacgcaatttttgacaagcacgaagttcaggagattaaggagaagatcctcaattccgattatgacgttgaggacttcttcgaaggtgagttttttaatttcgtgctcactcaagagggtatcgacgtgtataatgcgatcatcggtgggttcgtgactgagtccggtgaaaagattaagggattgaacgagtatatcaacctttacaaccaaaagacgaaacagaagctgccaaagttcaagcctctttacaaacaggttctttcagaccgcgagtcactctcgttctatggggagggctacacttcggatgaggaagtcctggaggtgttcaggaatactctcaataagaattcggagattttctcttctataaaaaaactggaaaagttgtttaagaattttgacgaatactctagcgccggcatatttgtgaaaaacggcccggccatatcaacgataagtaaagatatcttcggcgaatggaacgtgatcagagacaaatggaacgcggagtatgacgatattcacctgaagaagaaggctgtcgtaacggagaagtacgaggatgatcgcaggaaaagcttcaaaaagatcggaagtttcagcctggaacagttgcaggagtatgctgacgccgatcttagcgtcgtcgagaagttgaaggagataatcatccaaaaggtcgacgagatatataaagtctatggatcaagtgaaaaactgttcgacgccgacttcgttttggagaagtccctgaagaagaacgacgctgttgttgccattatgaaggatctgctcgacagcgtgaagagtttcgagaactatattaaggcttttttcggggaggggaaggagactaacagagatgagtccttctacggagacttcgtcctcgcgtacgatatactccttaaggtagaccacatctacgacgcaatcagaaattacgtgacacaaaagccgtacagcaaggacaagttcaaactctacttccagaacccccagttcatgggcggctgggacaaggacaaggaaacggattacagggctacgatcctgaggtatggttcaaaatactacttggcgattatggacaagaagtacgccaagtgtctccagaagattgacaaagacgatgtcaatggcaattatgagaagatcaactacaagctgcttccgggtccgaacaagatgctcccaaaggttttcttcagcaagaaatggatggcctactataacccaagcgaggacatccagaagatttataagaacggtacgttcaagaagggcgacatgttcaatcttaacgactgtcacaagctgatcgacttcttcaaagactcaattagccggtacccaaagtggtctaacgcctatgacttcaacttttcggaaaccgagaagtacaaggatatagccggattttatagagaggtggaagagcagggctacaaggtgtcattcgagtccgccagcaagaaggaagtggacaagctcgtggaagagggtaagctctacatgttccagatttataataaagactttagcgataagagccacgggacacctaatctccacacaatgtatttcaagctgctcttcgacgagaataaccacggccaaatcaggttgtcaggaggggctgaactcttcatgcggcgcgctagccttaagaaggaggagcttgtagtccaccctgcgaatagtccaattgcgaataagaacccggacaatcctaaaaagactacaacattgagctacgacgtgtacaaggataagaggttttccgaggatcagtac**GA*A*CTC**cacatcccgattgcgatcaacaagtgcccaaagaatattttcaagataaacacagaggtgcgtgtactcctgaagcatgacgacaatccttacgtcattgggattgatcggggcgagaggaacctcctctatattgtggtggtggacgggaaggggaacatagtcgaacagtactcccttaacgaaataattaacaatttcaacggcatccgtatcaagaccgactaccattcgttgctggacaagaaggagaaggagagatttgaggcgcggcaaaattggacaagtatcgagaacatcaaggaactcaaagcaggttatatctctcaagttgtgcataagatatgcgagctggttgagaagtatgacgcagtgatcgctcttgaggacctcaactcgggctttaagaattctagagttaaagtggagaagcaggtctatcaaaagttcgagaagatgcttatagataagctcaactacatggtcgataagaaatcgaacccatgtgccaccggcggcgcactcaaaggttaccaaataacaaacaaattcgagtccttcaaatcgatgagtactcagaatgggttcatattttatataccggcgtggcttacgtctaagatcgacccgtcaactggttttgtcaacctgttgaagacgaaatacacgtccattgccgattcgaaaaagttcatatctagttttgatcgtattatgtacgtcccagaggaagatcttttcgagtttgctctcgactacaaaaacttttcgcggaccgatgcggattacattaaaaaatggaaactctattcgtacggcaacagaatcaggatttttcgcaaccctaagaagaataacgtctttgattgggaggaagtttgcttgactagcgcgtacaag**GAGCT*G***tttaataagtatggcattaactaccaacagggtgatatcagagcactgctttgcgaacaatctgacaaggctttctactcatccttcatggctttgatgagcctgatgctccagatgagaaattcaattacaggcagaaccgacgtggatttcttgatctccccggttaaaaattctgatggcatcttttacgatagcaggaactatgaagcgcaagagaatgcgattctgccaaaaaatgcagacgccaacggtgcctataacatcgccaggaaagtcctgtgggcgatcggccagttcaaaaaggccgaagacgaaaaattggacaaggtcaaaatcgctatcagcaacaaagagtggctggagtatgctcagacatccgtaaagcataagcgtcctgctgccaccaaaaaggccggacaggctaagaaaaagaagtgaGAGCTCagagctttcgttcgtatcatcggtttcgacaacgttcgtcaagttcaatgcatcagtttcattgcgcacacaccagaatcctactgagtttgagtattatggcattgggaaaactgtttttcttgtaccatttgttgtgcttgtaatttactgtgttttttattcggttttcgctatcgaactgtgaaatggaaatggatggagaagagttaatgaatgatatggtccttttgttcattctcaaattaatattatttgttttttctcttatttgttgtgtgttgaatttgaaattataagagatatgcaaacattttgttttgagtaaaaatgtgtcaaatcgtggcctctaatgaccgaagttaatatgaggagtaaaacacttgtagttgtaccattatgcttattcactaggcaacaaatatattttcagacctagaaaagctgcaaatgttactgaatacaagtatgtcctcttgtgttttagacatttatgaactttcctttatgtaattttccagaatccttgtcagattctaatcattgctttataattatagttatactcatggatttgtagttgagtatgaaaatattttttaatgcattttatgacttgccaattgattgacaacgaattcgtaatcatggtcatagctgtttcctgtgtgaaattgttatccgctcacaattccacacaacatacgagccggaagcataaagtgtaaagcctggggtgcctaatgagtgagctaactcacattaattgcgttgcgctcactgcccgctttccagtcgggaaacctgtcgtgccagctgcattaatgaatcggccaacgcgcggggagaggcggtttgcgtattggctagagcagcttgccaacatggtggagcacgacactctcgtctactccaagaatatcaaagatacagtctcagaagaccaaagggctattgagacttttcaacaaagggtaatatcgggaaacctcctcggattccattgcccagctatctgtcacttcatcaaaaggacagtagaaaaggaaggtggcacctacaaatgccatcattgcgataaaggaaaggctatcgttcaagatgcctctgccgacagtggtcccaaagatggacccccacccacgaggagcatcgtggaaaaagaagacgttccaaccacgtcttcaaagcaagtggattgatgtgaacatggtggagcacgacactctcgtctactccaagaatatcaaagatacagtctcagaagaccaaagggctattgagacttttcaacaaagggtaatatcgggaaacctcctcggattccattgcccagctatctgtcacttcatcaaaaggacagtagaaaaggaaggtggcacctacaaatgccatcattgcgataaaggaaaggctatcgttcaagatgcctctgccgacagtggtcccaaagatggacccccacccacgaggagcatcgtggaaaaagaagacgttccaaccacgtcttcaaagcaagtggattgatgtgatatctccactgacgtaagggatgacgcacaatcccactatccttcgcaagaccCttcctctatataaggaagttcatttcatttggagaggacacgctgaaatcaccagtctctctctacaaatctatctctctcgagctttcgcagatctgtcgatcgaccatggggattgaacaagatggattgcacgcaggttctccggccgcttgggtggagaggctattcggctatgactgggcacaacagacaatcggctgctctgatgccgccgtgttccggctgtcagcgcaggggcgcccggttctttttgtcaagaccgacctgtccggtgccctgaatgaactccaggacgaggcagcgcggctatcgtggctggccacgacgggcgttccttgcgcagctgtgctcgacgttgtcactgaagcgggaagggactggctgctattgggcgaagtgccggggcaggatctcctgtcatctcaccttgctcctgccgagaaagtatccatcatggctgatgcaatgcggcggctgcatacgcttgatccggctacctgcccattcgaccaccaagcgaaacatcgcatcgagcgagcacgtactcggatggaagccggtcttgtcgatcaggatgatctggacgaagagcatcaggggctcgcgccagccgaactgttcgccaggctcaaggcgcgcatgcccgacggcgaggatctcgtcgtgacacatggcgatgcctgcttgccgaatatcatggtggaaaatggccgcttttctggattcatcgactgtggccggctgggtgtggcggaccgctatcaggacatagcgttggctacccgtgatattgctgaagagcttggcggcgaatgggctgaccgcttcctcgtgctttacggtatcgccgctcccgattcgcagcgcatcgccttctatcgccttcttgacgagttcttctgagcgggactctggggttcggatcgatcctctagctagagtcgatcgacaagctcgagtttctccataataatgtgtgagtagttcccagataagggaattagggttcctatagggtttcgctcatgtgttgagcatataagaaacccttagtatgtatttgtatttgtaaaatacttctatcaataaaatttctaattcctaaaaccaaaatccagtactaaaatccagatcccccgaattaattcggcgttaattcagtacattaaaaacgtccgcaatgtgttattaagttgtctaagcgtcaatttgtttacaccacaatatatcctgccaccagccagccaacagctccccgaccggcagctcggcacaaaatcaccactcgatacaggcagcccatcagtccgggacggcgtcagcgggagagccgttgtaaggcggcagactttgctcatgttaccgatgctattcggaagaacggcaactaagctgccgggtttgaaacacggatgatctcgcggagggtagcatgttgattgtaacgatgacagagcgttgctgcctgtgatcaccgcggtttcaaaatcggctccgtcgatactatgttatacgccaactttgaaaacaactttgaaaaagctgttttctggtatttaaggttttagaatgcaaggaacagtgaattggagttcgtcttgttataattagcttcttggggtatctttaaatactgtagaaaagaggaaggaaataataaatggctaaaatgagaatatcaccggaattgaaaaaactgatcgaaaaataccgctgcgtaaaagatacggaaggaatgtctcctgctaaggtatataagctggtgggagaaaatgaaaacctatatttaaaaatgacggacagccggtataaagggaccacctatgatgtggaacgggaaaaggacatgatgctatggctggaaggaaagctgcctgttccaaaggtcctgcactttgaacggcatgatggctggagcaatctgctcatgagtgaggccgatggcgtcctttgctcggaagagtatgaagatgaacaaagccctgaaaagattatcgagctgtatgcggagtgcatcaggctctttcactccatcgacatatcggattgtccctatacgaatagcttagacagccgcttagccgaattggattacttactgaataacgatctggccgatgtggattgcgaaaactgggaagaagacactccatttaaagatccgcgcgagctgtatgattttttaaagacggaaaagcccgaagaggaacttgtcttttcccacggcgacctgggagacagcaacatctttgtgaaagatggcaaagtaagtggctttattgatcttgggagaagcggcagggcggacaagtggtatgacattgccttctgcgtccggtcgatcagggaggatatcggggaagaacagtatgtcgagctattttttgacttactggggatcaagcctgattgggagaaaataaaatattatattttactggatgaattgttttagtacctagaatgcatgaccaaaatcccttaacgtgagttttcgttccactgagcgtcagaccccgtagaaaagatcaaaggatcttcttgagatcctttttttctgcgcgtaatctgctgcttgcaaacaaaaaaaccaccgctaccagcggtggtttgtttgccggatcaagagctaccaactctttttccgaaggtaactggcttcagcagagcgcagataccaaatactgtccttctagtgtagccgtagttaggccaccacttcaagaactctgtagcaccgcctacatacctcgctctgctaatcctgttaccagtggctgctgccagtggcgataagtcgtgtcttaccgggttggactcaagacgatagttaccggataaggcgcagcggtcgggctgaacggggggttcgtgcacacagcccagcttggagcgaacgacctacaccgaactgagatacctacagcgtgagctatgagaaagcgccacgcttcccgaagggagaaaggcggacaggtatccggtaagcggcagggtcggaacaggagagcgcacgagggagcttccagggggaaacgcctggtatctttatagtcctgtcgggtttcgccacctctgacttgagcgtcgatttttgtgatgctcgtcaggggggcggagcctatggaaaaacgccagcaacgcggcctttttacggttcctggccttttgctggccttttgctcacatgttctttcctgcgttatcccctgattctgtggataaccgtattaccgcctttgagtgagctgataccgctcgccgcagccgaacgaccgagcgcagcgagtcagtgagcgaggaagcggaagagcgcctgatgcggtattttctccttacgcatctgtgcggtatttcacaccgcatatggtgcactctcagtacaatctgctctgatgccgcatagttaagccagtatacactccgctatcgctacgtgactgggtcatggctgcgccccgacacccgccaacacccgctgacgcgccctgacgggcttgtctgctcccggcatccgcttacagacaagctgtgaccgtctccgggagctgcatgtgtcagaggttttcaccgtcatcaccgaaacgcgcgaggcagggtgccttgatgtgggcgccggcggtcgagtggcgacggcgcggcttgtccgcgccctggtagattgcctggccgtaggccagccatttttgagcggccagcggccgcgataggccgacgcgaagcggcggggcgtagggagcgcagcgaccgaagggtaggcgctttttgcagctcttcggctgtgcgctggccagacagttatgcacaggccaggcgggttttaagagttttaataagttttaaagagttttaggcggaaaaatcgccttttttctcttttatatcagtcacttacatgtgtgaccggttcccaatgtacggctttgggttcccaatgtacgggttccggttcccaatgtacggctttgggttcccaatgtacgtgctatccacaggaaacagaccttttcgacctttttcccctgctagggcaatttgccctagcatctgctccgtacattaggaaccggcggatgcttcgccctcgatcaggttgcggtagcgcatgactaggatcgggccagcctgccccgcctcctccttcaaatcgtactccggcaggtcatttgacccgatcagcttgcgcacggtgaaacagaacttcttgaactctccggcgctgccactgcgttcgtagatcgtcttgaacaaccatctggcttctgccttgcctgcggcgcggcgtgccaggcggtagagaaaacggccgatgccgggatcgatcaaaaagtaatcggggtgaaccgtcagcacgtccgggttcttgccttctgtgatctcgcggtacatccaatcagctagctcgatctcgatgtactccggccgcccggtttcgctctttacgatcttgtagcggctaatcaaggcttcaccctcggataccgtcaccaggcggccgttcttggccttcttcgtacgctgcatggcaacgtgcgtggtgtttaaccgaatgcaggtttctaccaggtcgtctttctgctttccgccatcggctcgccggcagaacttgagtacgtccgcaacgtgtggacggaacacgcggccgggcttgtctcccttcccttcccggtatcggttcatggattcggttagatgggaaaccgccatcagtaccaggtcgtaatcccacacactggccatgccggccggccctgcggaaacctctacgtgcccgtctggaagctcgtagcggatcacctcgccagctcgtcggtcacgcttcgacagacggaaaacggccacgtccatgatgctgcgactatcgcgggtgcccacgtcatagagcatcggaacgaaaaaatctggttgctcgtcgcccttgggcggcttcctaatcgacggcgcaccggctgccggcggttgccgggattctttgcggattcgatcagcggccgcttgccacgattcaccggggcgtgcttctgcctcgatgcgttgccgctgggcggcctgcgcggccttcaacttctccaccaggtcatcacccagcgccgcgccgatttgtaccgggccggatggtttgcgaccgctcacgccgattcctcgggcttgggggttccagtgccattgcagggccggcagacaacccagccgcttacgcctggccaaccgcccgttcctccacacatggggcattccacggcgtcggtgcctggttgttcttgattttccatgccgcctcctttagccgctaaaattcatctactcatttattcatttgctcatttactctggtagctgcgcgatgtattcagatagcagctcggtaatggtcttgccttggcgtaccgcgtacatcttcagcttggtgtgatcctccgccggcaactgaaagttgacccgcttcatggctggcgtgtctgccaggctggccaacgttgcagccttgctgctgcgtgcgctcggacggccggcacttagcgtgtttgtgcttttgctcattttctctttacctcattaactcaaatgagttttgatttaatttcagcggccagcgcctggacctcgcgggcagcgtcgccctcgggttctgattcaagaacggttgtgccggcggcggcagtgcctgggtagctcacgcgctgcgtgatacgggactcaagaatgggcagctcgtacccggccagcgcctcggcaacctcaccgccgatgcgcgtgcctttgatcgcccgcgacacgacaaaggccgcttgtagccttccatccgtgacctcaatgcgctgcttaaccagctccaccaggtcggcggtggcccatatgtcgtaagggcttggctgcaccggaatcagcacgaagtcggctgccttgatcgcggacacagccaagtccgccgcctggggcgctccgtcgatcactacgaagtcgcgccggccgatggccttcacgtcgcggtcaatcgtcgggcggtcgatgccgacaacggttagcggttgatcttcccgcacggccgcccaatcgcgggcactgccctggggatcggaatcgactaacagaacatcggccccggcgagttgcagggcgcgggctagatgggttgcgatggtcgtcttgcctgacccgcctttctggttaagtacagcgataaccttcatgcgttccccttgcgtatttgtttatttactcatcgcatcatatacgcagcgaccgcatgacgcaagctgttttactcaaatacacatcacctttttagacggcggcgctcggtttcttcagcggccaagctggccggccaggccgccagcttggcatcagacaaaccggccaggatttcatgcagccgcacggttgagacgtgcgcgggcggctcgaacacgtacccggccgcgatcatctccgcctcgatctcttcggtaatgaaaaacggttcgtcctggccgtcctggtgcggtttcatgcttgttcctcttggcgttcattctcggcggccgccagggcgtcggcctcggtcaatgcgtcctcacggaaggcaccgcgccgcctggcctcggtgggcgtcacttcctcgctgcgctcaagtgcgcggtacagggtcgagcgatgcacgccaagcagtgcagccgcctctttcacggtgcggccttcctggtcgatcagctcgcgggcgtgcgcgatctgtgccggggtgagggtagggcgggggccaaacttcacgcctcgggccttggcggcctcgcgcccgctccgggtgcggtcgatgattagggaacgctcgaactcggcaatgccggcgaacacggtcaacaccatgcggccggccggcgtggtggtgtcggcccacggctctgccaggctacgcaggcccgcgccggcctcctggatgcgctcggcaatgtccagtaggtcgcgggtgctgcgggccaggcggtctagcctggtcactgtcacaacgtcgccagggcgtaggtggtcaagcatcctggccagctccgggcggtcgcgcctggtgccggtgatcttctcggaaaacagcttggtgcagccggccgcgtgcagttcggcccgttggttggtcaagtcctggtcgtcggtgctgacgcgggcatagcccagcaggccagcggcggcgctcttgttcatggcgtaatgtctccggttctagtcgcaagtattctactttatgcgactaaaacacgcgacaagaaaacgccaggaaaagggcagggcggcagcctgtcgcgtaacttaggacttgtgcgacatgtcgttttcagaagacggctgcactgaacgtcagaagccgactgcactatagcagcggaggggttggatcaaagtactttgatcccgaggggaaccctgtggttggcatgcacatacaaatggacgaacggataaaccttttcacgcccttttaaatatccgttattctaa

**II. Schematic map with key elements in T-DNA.** Except poly(T) and Cas12a, the other elements are labeled as described by (Xing et al. 2014).

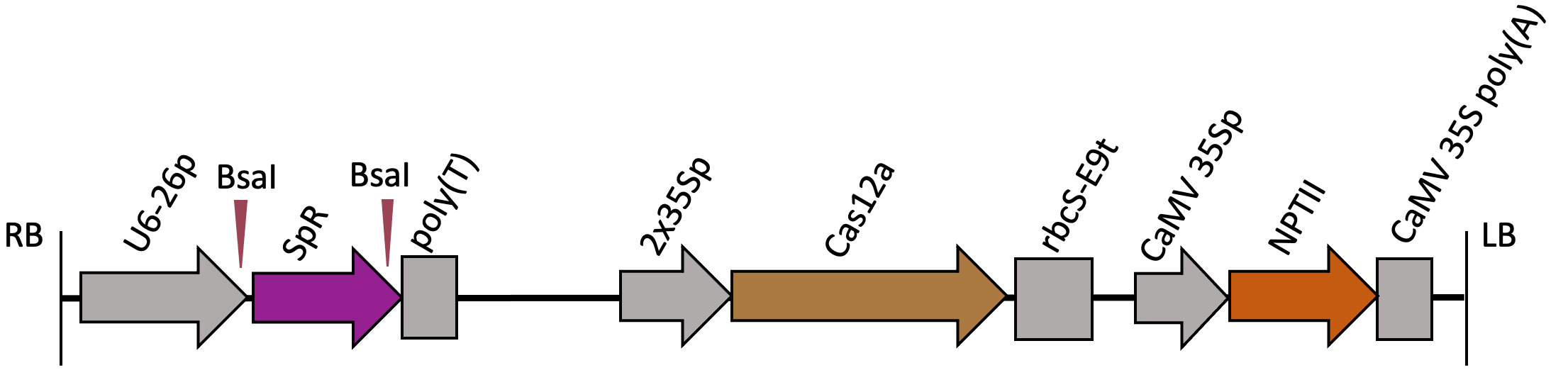

For cloning guide RNAs to pKSE401-Cas12a using BsaI sites to replace SpR, the oligos should have overhangs as depicted below.

F oligos (5’-3’): 5’-ATTGNNNNNNNNN---NNNN-3’

R oligos (3’-5’): 3’-NNNNNNNNN---NNNNAAAA-5’
