## Supplementary material for "Genome editing of papaya using both Cas9 and Cas12a": Table S1

**Table S1. Target sequences of guide RNAs and primers used to amplify the fragments for *in vitro* cleavage activity assays and mutation detection.**

| Target gene-Cas | Guide RNA ID | Primer sequences to amplify the fragment for in vitro cleavage assay | Primers used for detection of mutations |
| --- | --- | --- | --- |
| CpPDS-Cas9 | sg16 | CpPDS-InVitro_F1:  5'-CCAGATAGACATTACCCAGAATC-3' | CpPDS-InVitro_F1:  5'-CCAGATAGACATTACCCAGAATC-3' |
|  |  | CpPDS-InVitro_R1:  5'-CTATGTCCTGGAATGAACTTCAC-3' | CpPDS-sg16mt_R1:  5'-GGATTCACTAACCCTAAATGC-3' |
|  | sg379 | CpPDS-InVitro_F2:  5'-GTAAGTGTACTCCTCAGGCCAGTG-3' | CpPDS_sg379mt-F1:  5'-GTTGTTGATTGCCTTATTCCCA-3' |
|  |  | CpPDS-InVitro_R2:  5'-GTGAACCTCAAATCTGCAGAAGC-3' | CpPDS_sg379mt-R1:  5'-GCGAAAGGCAAGCACAA-3' |
| CpPDS-Cas12a | cr253 | As above for sg16 | CpPDS-sg16_F1:  5’-GGTGTTTCTGCGGCGAGCTT-3’ |
|  |  |  | CpPDS-Exon2_R1:  5’-CGATCTTCAATGGTCTCGATGGACG-3’ |
|  | cr423rc | As above for sg379 | As above for sg379 |
| CpMLO6-Cas9 | sg254 | CpMLO6-InVitro_F1:  5'-CCTTCATATGTCCGTATCACTG-3' | CpMLO6-sg254mt_F1:  5'-GAGAGCTCTGTACGAATCACTTG-3' |
|  |  | CpMLO6-InVirto_R1:  5'-ACATCCATACGCTACGTACTTC-3' | CpMLO6-sg254mt_R1:  5'-GCACATTTATCAGTAGAGGCA-3' |
|  | sg439 | CpMLO6-InVitro_F2:  5'-GAAGTACGTAGCGTATGGATGT-3' | CpMLO6-sg439mt_F1:  5'-GGACTGGATGTGTCTTATAAACGC-3' |
|  |  | CpMLO6-InVitro_R2:  5'-CGGCCAGGATTCTCTATTGAG-3' | CpMLO6-sg439mt_R1:  5'-CTTAGCTCTGCCCAAAGCC-3' |
| CpMLO6-Cas12a | cr292rc | As above for sg254 | As above for sg254 |
|  | cr535 | As above for sg439 | CpMLO6-sg439mt_F1:  5'-GGACTGGATGTGTCTTATAAACGC-3' |
|  |  |  | CpMLO6-Sg535mt-R1:  5'-GTTCAGACCAACAGGGTCCAGATC-3' |
