## Supplementary material for "Genome editing of papaya using both Cas9 and Cas12a": Table S2

**Table S2. Oligos used for constructing the plasmids for genome editing of *CpPDS* and *CpMLO6*.** The sequences and reverse complementary sequences of the targets of guide RNAs are underlined. The full-length direct repeats (FLDR) of CRISPR-Cas12a are shaded.

| **pKSE401::CpPDS** | CpPDS-sg379-BsF: 5’-ATATATGGTCTCGATT**GCAAAATATTTGGCAGATGC**GTT-3’ |
| --- | --- |
|  | CpPDS-sg379-F0: 5’-T**GCAAAATATTTGGCAGATGC**GTTTTAGAGCTAGAAATAGC-3’ |
|  | CpPDS-sg16-R0: 5’-AAC**AAGCTCGCCGCAGAAACAC**CAATCTCTTAGTCGACTCTAC-3’ |
|  | CpPDS-sg16-BsR:5’-ATTATTGGTCTCGAAAC**AAGCTCGCCGCAGAAACAC**CAA-3’ |
| **pKSE401::CpMLO6** | CpMLO6-sg439-BsF:5’-ATATATGGTCTCGATT**GTCCCATTTGTATCCGAAGA**GTT-3’ |
|  | CpMLO6-sg439-F0: 5’-T**GTCCCATTTGTATCCGAAGA**GTTTTAGAGCTAGAAATAGC-3’ |
|  | CpMLO6-sg254-R0: 5’-AAC**CGTGGCTCCAATTGACTTC**CAATCTCTTAGTCGACTCTAC-3’ |
|  | CpMLO6-sg254-BsR: 5’-ATTATTGGTCTCGAAAC**CGTGGCTCCAATTGACTTC**CAA-3’ |
| **pKSE401-Cas12a::CpPDS** | Cas12a-CpPDs-F:  5’-attgTTTCAAAGATTAAATAATTTCTACTAAGTGTAGAT**CACCTAGAACATCTCTTG** CTTTTCAAAGATTAAATAATTTCTACTAAGTGTAGAT**TGTAGATTACCCGAGACCAG**TTTCAAAGATTAAATAATTTCTACTAAGTGTAGAT-3’  Cas12a-CpPDs-R:  5’-aaaaATCTACACTTAGTAGAAATTATTTAATCTTTGAAA**CTGGTCTCGGGTAATCTA** *CA*ATCTACACTTAGTAGAAATTATTTAATCTTTGAAA**AGCAAGAGATGTTCTAGGTG**ATCTACACTTAGTAGAAATTATTTAATCTTTGAAA-3’ |
| **pKSE401-Cas12a::CpMLO6-cr292rc** | cr292rc-F:  5’-attgTTTCAAAGATTAAATAATTTCTACTAAGTGTAGAT**TCCGATTCCTCCTCTTGCT** **T**TTTCAAAGATTAAATAATTTCTACTAAGTGTAGAT-3’  cr292rc-R:  5’-aaaaATCTACACTTAGTAGAAATTATTTAATCTTTGAAA**AAGCAAGAGGAGGAATC** **GGA**ATCTACACTTAGTAGAAATTATTTAATCTTTGAAA-3’ |
| **pKSE401-Cas12a::CpMLO6-cr535** | cr535-F: 5’-attgTTTCAAAGATTAAATAATTTCTACTAAGTGTAGAT **CTGTATTATCACCATGGCT T**TTTCAAAGATTAAATAATTTCTACTAAGTGTAGAT  Cr535-R;  5’-aaaaATCTACACTTAGTAGAAATTATTTAATCTTTGAAA**AAGCCATGGTGATAATACA** **G**ATCTACACTTAGTAGAAATTATTTAATCTTTGAAA |
